# Lamellipodia-like actin networks in cells lacking WAVE Regulatory Complex

**DOI:** 10.1101/2021.06.18.449030

**Authors:** Frieda Kage, Hermann Döring, Magdalena Mietkowska, Matthias Schaks, Franziska Grüner, Stephanie Stahnke, Anika Steffen, Mathias Müsken, Theresia E. B. Stradal, Klemens Rottner

**Author notes:** Author for correspondence: Klemens Rottner.

## Abstract

Cell migration frequently involves the formation of lamellipodia induced by Rac GTPases mediating activation of WAVE Regulatory Complex (WRC) driving Arp2/3 complex-dependent actin assembly. Previous genome editing studies solidified the view of an essential, linear pathway employing aforementioned components. Using disruption of the WRC subunit Nap1 and its paralogue Hem1 followed by serum and growth factor stimulation or expression of active GTPases now revealed a pathway to formation of Arp2/3 complex-dependent, lamellipodia-like structures (LLS) that require both Rac and Cdc42, but not WRC. These observations were independent of WRC subunit eliminated and coincided with the lack of recruitment of Ena/VASP family actin polymerases. Moreover, aside from the latter, induced LLS contained all common lamellipodial regulators tested, including cortactin, the Ena/VASP ligand lamellipodin or FMNL subfamily formins. Our studies thus establish the existence of a signaling axis to Arp2/3 complex-dependent actin remodeling at the cell periphery operating without WRC and Ena/VASP.

## Introduction

Protrusions at the cell periphery are frequently driven by actin networks generated through the actin filament branching activity of Arp2/3 complex (Rotty et al., 2013). With the exception of bundles containing parallel actin filaments commonly referred to as filopodia and likely driven by the Ena/VASP family of actin polymerases, formins or both (Gallop, 2020; Rottner et al., 2017), these actin networks give rise to various structures in distinct cell types. These include lamellipodia and ruffles mediating migration or perhaps macropinocytosis, the phagocytic cups of professional phagocytes or the actin-dependent protuberances accompanying the induced entry of specific pathogens (Krause and Gautreau, 2014; Stradal and Schelhaas, 2018).

Actin remodeling in mammalian cells is largely regulated by small GTPases of the Rho family (Aspenstrom et al., 2004), and members of the Rac subfamily (Rac1, 2 and 3) have emerged as key GTPases obligatory for membrane ruffling and lamellipodia formation in cell types ranging from melanoma to platelets (McCarty et al., 2005; Schaks et al., 2018; Steffen et al., 2013). In early studies, there were conflicting ideas about how precisely Rac connects to WAVE, the key activator of Arp2/3 complex in membrane ruffles and lamellipodia (Machesky and Insall, 1998). The most promising, initial candidate was IRSp53 (Miki et al., 2000), which was also found to co-localize with WAVE at the edges of protruding lamellipodia (Nakagawa et al., 2003). However, even earlier studies had already described an alternative Rac interactor, termed Sra1 (Specifically Rac-associated protein) (Eden et al., 2002; Kobayashi et al., 1998), which is now well established to constitute even two interaction surfaces with Rac on heteropentameric WRC (Chen et al., 2017). Early studies using RNA interference in mammalian and *Drosophila* cells indicated that all WRC subunits are essential for Rac-mediated actin remodeling (Innocenti et al., 2004; Kunda et al., 2003; Rogers et al., 2003; Steffen et al., 2004). Genetic deletion of WRC subunits in both mammalian and *Dictyostelium* cells have hitherto confirmed the key role of WRC in Rac-mediated actin remodeling (Schaks et al., 2018; Whitelaw et al., 2020), albeit with phenotypes obtained upon targeting distinct WRC subunits in amoeba being admittedly more heterogeneous (Bear et al., 1998; Davidson and Insall, 2013; Litschko et al., 2017). Based on all these results, WRC was considered to be crucial, if not obligatory, for Arp2/3 complex-dependent actin remodeling in cell edge protrusions, i.e. downstream of Rac (Rottner and Schaks, 2019), although some doubts have remained. As examples, *Dictyostelium* cells lacking Scar were reported to engage WASP instead of Scar, although this required increased Rac activities in these cells (Veltman et al., 2012), and *C. elegans* neuroblasts appeared to employ both WAVE and WASP for leading edge Arp2/3 activation (Zhu et al., 2016). Furthermore, in spite of a lack of phenotype concerning the efficiency of membrane ruffling and lamellipodia protrusion in different mammalian cell types lacking N-WASP or the mostly hematopoietic WASP (Leithner et al., 2016; Lommel et al., 2001; Snapper et al., 2001), mature dendritic cells displayed WASP accumulation at the cell periphery if forced to protrude under confinement. Likewise, WRC loss of function in tumor cells migrating in 3D appeared to promote N-WASP-dependent invasion, indicating complex, perhaps condition- and context-dependent interrelationships between WRC and WASP/N-WASP functions in regulating Arp2/3—dependent actin remodeling at the cell periphery.

In this study, we establish and characterize a hitherto unrecognized lamellipodial-like activity in mouse melanoma and fibroblast cells lacking WRC, but involving various lamellipodial regulators aside from Ena/VASP family proteins as only exception.

## Results and discussion

### Nap1 null clones display distinct lamellipodial phenotypes due to differential Hem1 expression

To obtain B16-F1 melanoma cell clones abolished for WRC function alternative to Sra1/PIR121- or WAVE KOs (Schaks et al., 2018; Tang et al., 2020), we initially disrupted the *Nckap1* gene giving rise to expression of Nap1 (Fig. S1A), hitherto thought to be essential for WRC function in this cell type (Steffen et al., 2006; Steffen et al., 2004). A subfraction of these clones (#6 and #21) were previously used as tools for cells with low lamellipodial Arp2/3 complex activities (Dolati et al., 2018), but not characterized in further detail. Clone #16 was also employed recently to show that cells lacking apparent lamellipodia display a significant reduction of turnover of peripheral actin filaments (see also below, Whitelaw et al., 2020), as expected (Steffen et al., 2014). Except for clones #1 and #2 showing strongly reduced but not abolished Nap1 expression and thus not used further, eleven additional clones lacked apparent Nap1 expression (Fig. S1A), five of which were confirmed to lack *Nckap1* wt alleles (Fig. S1B) and thus analyzed further. Indeed, two of these single cell clones (#16 and #23) were virtually devoid of lamellipodia (Fig. 1A), whereas #6 occasionally and #21 quite frequently displayed cells with lamellipodia, which certainly came as surprise (see also Dolati et al., 2018). These phenotypic observations also correlated with modest but clearly detectable expression of Sra1/PIR121 and WAVE in extracts from bulk populations of clone #21 cells, but not the others (Figs. 1B, S1C). However, rare exception of individual cells with underdeveloped, very narrow lamellipodia could even be found in the remaining clones (for superresolution images see Fig. S2). Expression levels of Abi and HSPC300 were also modestly reduced in these clones, but were likely not limiting (Figs. 1B, S1C). Further characterization of lamellipodia formed in clone #21 cells confirmed the presence - aside from Arp2/3 complex - of all WRC subunits except for Nap1 (Fig. 1C), suggesting that functionality of these WRCs was achieved through expression of the mostly hematopoietic Hem1 (Park et al., 2010). And indeed, despite failure to detect Hem1 in regular cell extracts of Nap1 KO#21, we were able to co-IP the protein with EGFP-tagged Sra1, just like WAVE or Abi, but only in clone #21, and not in B16-F1 wildtype (Fig. S1D). These data strongly suggested compensatory Hem1 expression in clone #21, which could even be enhanced upon subcloning, giving rise to Nap1 KO #21-12, displaying WAVE expression and lamellipodia formation efficiency indistinguishable from B16-F1 wildtype (Fig. 1D). To prove Hem1 expression in this clone, we CRISPR/Cas9-engineered EGFP into the *hem1*-gene locus, and confirmed EGFP-Hem1 expression in three independent clones by Western and its accumulation at lamellipodia tips in single cell fluorescence microscopy (Fig. 1E).

**Figure 1.**
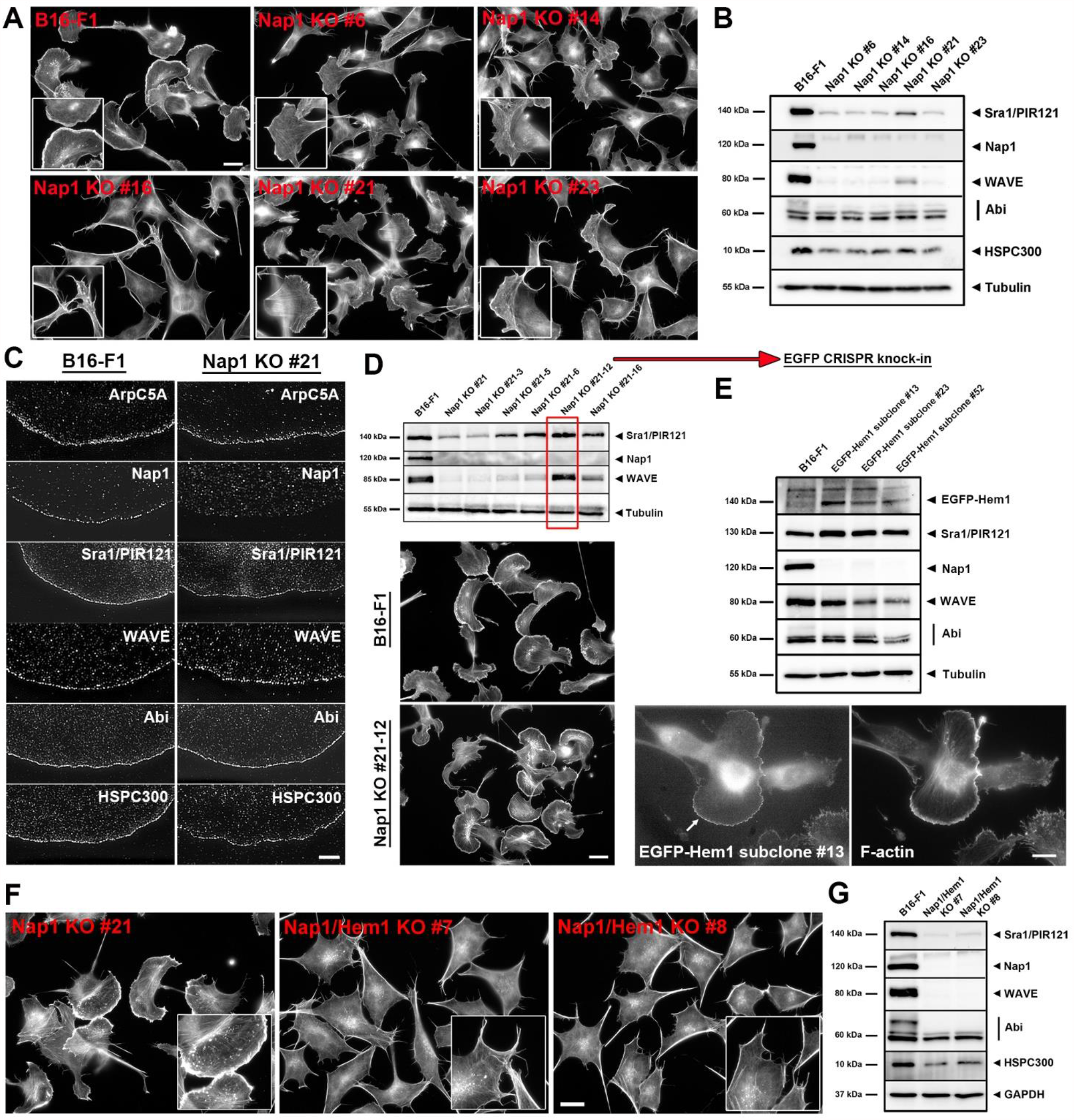
Elimination of canonical lamellipodia in B16-F1 cells requires removal of both Nap1 and Hem1. **(A)** Comparison of actin cytoskeleton morphologies of B16-F1 wildtype and isolated Nap1 single KO clones (see Fig. S1) stained with phalloidin. Note that Nap1 KO in B16-F1 cells altered lamellipodia frequency and morphology to variable extents in distinct clones, but did not completely abrogate their formation (insets). Scale bar, 20 µm. **(B)** Western blotting of whole cell extracts of selected Nap1 KO clones and B16-F1 cells as control to detect expression levels of other WRC subunits, as indicated. α-Tubulin served as loading control. **(C)** Representative SIM-images of lamellipodial sections from either B16-F1 cells or Nap1 KO #21. Antibody staining demonstrates accumulation of Arp2/3 complex (subunit ArpC5A) and all WRC subunits besides Nap1 in lamellipodia formed by Nap1 KO clone #21. Scale bar, 5 µm. Subcloning of Nap1 KO clone #21 in order to enrich for a cell population with high Hem1 expression. Despite the lack of Nap1, derived subclone #21-12 (red box) was found to possess expression levels of WRC subunits identical to parental B16-F1 cells, indicative of further increased, compensatory Hem1 expression. Accordingly, cellular morphologies and lamellipodia formation were indistinguishable from B16-F1 wildtype controls (bottom), as shown by phalloidin staining. Scale bar, 20 µm. **(E)** EGFP-insertion upstream of the *hem1* gene locus to give rise to a fusion protein proving robust, endogenous Hem1 protein expression in subclone #21-12 using a CRISPR-mediated knock-in approach (red arrow, see Materials and Methods). Expression of EGFP-tagged Hem1 at its expected molecular weight in three, independently isolated knock-in clones as verified by Western blotting using anti-EGFP antibodies. Bottom: Cell images displaying the formation of lamellipodia stained with phalloidin (right) and tipped by EGFP-Hem1 (left, white arrow); representative data shown for Nap1 KO #21-12 + EGFP-Hem1 subclone #13. Scale bar, 10 µm. **(F)** Overview images of phalloidin-stained Nap1 KO clone #21 and two clones derived from the latter by consecutive, additional CRISPR/Cas9-mediated genetic disruption of *hem1* (see also Fig. S1 for more details). Note that additional removal of Hem1 leads to complete loss of lamellipodia formation (insets), at least in standard growth conditions on laminin. Scale bar, 20 µm. **(G)** Western blotting of whole cell extracts of B16-F1 wildtype *versus* Nap1/Hem1 double null cell lines, confirming still differential reduction of expression of distinct WRC subunits, yet virtually complete elimination of WAVE expression (as compared to Nap1 single KO clones shown in B; for quantitations see Fig. S1).

We also employed Nap1 KO clones #16 and #23 with largely lacking lamellipodial activity and hence endogenous Hem1 expression for comparative rescue experiments with EGFP-tagged Nap1 *versus* Hem1, to explore the possibility of differential activities between the two genes concerning lamellipodia formation or cell migration. Indeed, previous studies established a single point mutation abrogating Hem1 function, which was without effect at the corresponding sequence position in Nap1 (Salzer et al., 2020). However, extent and frequency of lamellipodia formation (Fig. S3A, B), WRC subunit expression (Fig. S3C) as well as migration efficiency (Fig. S3D–F) were all equally efficiently restored in Nap1 KOs with either EGFP-Nap1 or ‐Hem1. Even the turnover of both fusion proteins at lamellipodia tips was comparably slow (t_1/2_ of app. 30 sec, Fig. S3G, H).

All these data confirmed lack of lamellipodia formation in specific, individual Nap1 KO clones (i.e. clones #16 and #23) to derive from virtually absent Hem1 expression, and their differential formation in alternatively isolated clones or subclones to derive from upregulation of Hem1 considered to be equally active to Nap1 at least concerning the lamellipodial or migration parameters analyzed here.

### Thresholds of WRC activity determine frequency of lamellipodia formation and coinciding efficiency of cell migration

To obtain cells lacking canonical lamellipodia, we first disrupted the *hem1* locus in Nap1 null cells. For screening purposes in the absence of a highly sensitive Hem1-specific western blotting antibody, Nap1 KO #21 cells were chosen and derived Nap1/Hem1 nulls isolated using lamellipodial deficiency as parameter. After selection of double KO clones based on morphological parameters (Fig. 1F), two were subjected to sequencing of the *hem1* locus (#7 and #8, Fig. S1E). Both clones harbored at least three alleles, all leading to gene disruption except for one in-frame deletion of residue 21 in clone #7. Due to the absence of lamellipodia in these conditions, we concluded that deletion of the aspartate residue at position 21 of Hem1 generates a non-functional variant of the protein. Expression of EGFP-tagged Hem1ΔD21 in Nap1/Hem1 double KO cells confirmed this view, since lamellipodia were not rescued by the mutant, but readily observed in the same cells transfected with EGFP-Hem1 wildtype (Fig. S4). Nap1/Hem1 double KO cell lines were virtually devoid of both Sra1/PIR121 and WAVE, and reduced in expression of Abi and HSPC300 to app. half of wildtype levels (Figs. 1G, S1F).

Interestingly, careful quantitative analysis of random cell migration in the five Nap1 single KO clones selected as shown in Fig. 1 revealed a roughly comparable reduction of average migration rates to about 0.4 µm/min (Fig. 2A), except for Nap1 KO #21, and highly similar to previous observations with Sra1/PIR121 double KO cells (Schaks et al., 2018). However, categorization of individually analyzed cells into those with and without lamellipodia revealed that the two subsets of cells roughly comprised two groups with astonishingly comparable migration speeds, app. 1.5 µm/min with and roughly one third of that without lamellipodia (Fig. 2B). The difference in clone #21 as compared to the remaining Nap1 KO clones arose from the fact that the majority of clone #21 cells displayed lamellipodia and migrated at wildtype levels, whereas the ratios of rapidly *versus* slowly migrating cells was reversed in all other clones. Clone #16 constituted the extreme case, as for the latter, no single lamellipodia-forming cell was found out of 397 cells in this experiment. The data thus showed that the differences between clones were mostly reflected by the number of cells able to form lamellipodia in these migration conditions. Even in clones displaying very low overall migration rates, as nicely illustrated in migration trajectory plots (Fig. 2C), such as for instance in clone #23, those few cells capable of lamellipodia formation (n=9 out of 382 cells analyzed) migrated at levels virtually identical to the majority of cells migrating with lamellipodia in wildtype B16-F1 (Fig. 2B). We conclude that although lamellipodium protrusion efficiency appeared to correlate with average WRC levels expressed in clones #6 *versus* #21 previously (Dolati et al., 2018), these subtle, lamellipodia-specific differences did not translate into distinct effectiveness of overall migration rates. Therefore, at the individual cell level, the capability to form a canonical, WRC-dependent lamellipodium was decisive for effective migration, irrespective of clone origin, and likely dependent on reaching a given threshold sufficient to form a lamellipodium (Fig. 2). In Nap1/Hem1 double KO clones, migration was not entirely abolished, but average rates strongly suppressed to below 0.3 µm/min, consistent with the complete absence of canonical lamellipodia in all cells examined (n>415, Fig. 2D and data not shown).

**Figure 2.**
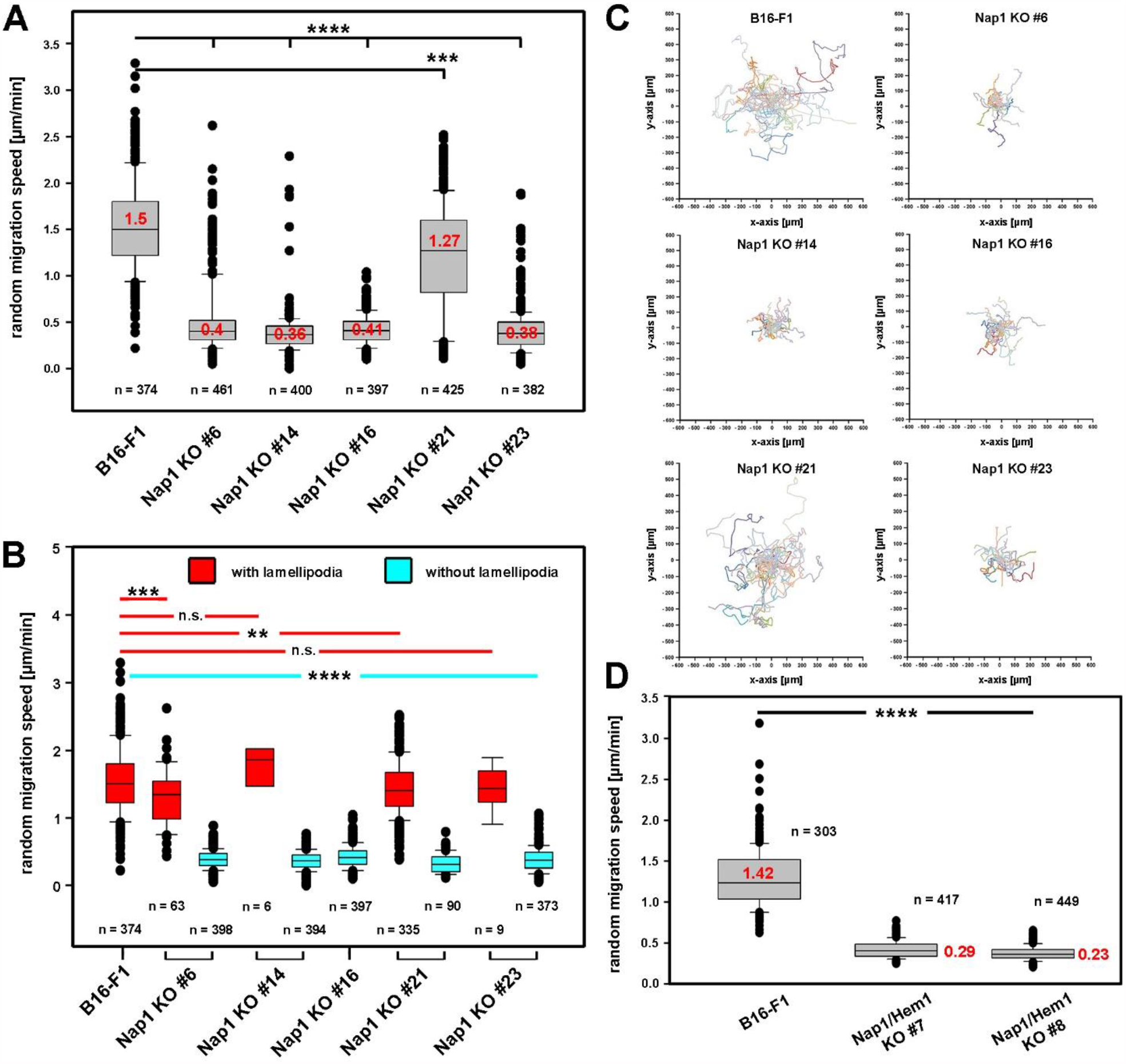
Effective migration of B16-F1 cells closely correlates with the capability of lamellipodia formation. **(A)** Random migration speed of Nap1 KO clones compared to parental B16-F1 cells. Box and whisker plots with boxes corresponding to 50% of data points (25%–75%), and whiskers corresponding to 80% (10%–90%). Outliers are shown as dots, red numbers in boxes correspond to medians. All Nap1 single KOs were markedly reduced in migration, except for KO #21, which was only moderately affected. Yet, all changes were found to be statistically significantly different. **(B)** Random migration data discriminating Nap1 KO cells with (red boxes) and without (blue boxes) lamellipodia show that lamellipodia-forming Nap1 KO cells migrate at rates indistinguishable from controls, while lamellipodia-deficient cells displayed an average migration speed of 0.4 µm/min. This level was highly reminiscent of lamellipodia-deficient cells lacking Sra1/PIR121 (Schaks et al., 2018). **(C)** Trajectory plots illustrating that the frequency of lamellipodia formation revealed in B correlates with overall migration efficiency. **(D)** Random migration speed of B16-F1 cells and in comparison, Nap1/Hem1 double KO cells. Note that disruption of both genes decreases migration speed even further to 0.26 µm/min on average. These data, together with previous observations on Sra1/PIR121 KO suggest additional, lamellipodia-independent functions of the Nap1/Hem1 module in migration. Statistics in all cases were done with one-way ANOVA with Dunnet’s adjustment for multiple comparisons; ^****^ = p ≤ 0.0001, ^***^ = p ≤ 0.001, ^**^ = p ≤ 0.01 and n.s. = p ≥ 0.05.

### Growth factor stimulation of B16-F1 cells can induce the formation of lamellipodia-like structures (LLS) devoid of WRC but dependent on Rac

Since more than two decades, B16-F1 melanoma cells constitute a well-established model system of actin-based protrusion, migrating by employing broad and flat lamellipodia in regular growth medium when seeded on laminin (Ballestrem et al., 1998; Rottner et al., 1999a; Svitkina et al., 2003). In these conditions, we have never seen any signs of lamellipodial networks formed at the periphery of cells lacking WRC subunits (see e.g. Fig. 1F and Salzer et al., 2020; Schaks et al., 2018).

However, along our efforts to challenge this model with various conditions stimulating actin remodeling, we found, to our surprise, that overnight starvation of Nap1/Hem1 double-null clones followed by treatment with growth factors in full medium triggered the formation of lamellipodia-like structures (LLS). Complementation of the growth medium with HGF, PDGF and EGF all gave comparable results, but we restricted ourselves to showing the data obtained with HGF only. Resulting structures appeared strongly reminiscent of canonical lamellipodia, albeit smaller in size as compared to those seen in B16-F1 wildtype (Fig. 3A). Next, and even more striking, we found that formation of these structures was independent of how WRC was inactivated, since Sra1/PIR121 KO cells were indistinguishable in this respect from Nap1/Hem1 KOs, and that they all contained Arp2/3 complex (for representative cell examples see Fig. 3B). In contrast, however, induction of these structures was only seen in WRC-deficient cells, but not in the absence of Rac1/2/3 (Fig. 3B). Line scan fluorescence quantifications of the cell periphery of stimulated cells revealed virtually identical peak accumulations for the lamellipodial markers Arp2/3 complex and cortactin, but again not in the absence of Rac-GTPases (Rac1/2/3). None of the genotypes displayed such peaks without stimulation (Fig. 3C). Next, we explored the accumulation of additional key lamellipodial regulators in these LLS, counterstaining as control either Arp2/3 complex or cortactin with mono- or polyclonal antibodies, respectively. The WRC subunit Abi1 was confirmed to be completely absent from these structures, as expected for cells lacking functional WRC (Fig. 4, top panel; see Fig. S5 top panels for representative images), although expression levels for this subunit were decreased to only about half of control cells (Fig. S1F). This suggested that WRC subunits cannot be recruited to the lamellipodium in the absence of the Rac-interacting Sra1/PIR121 module or its tight interactor Nap1/Hem1, the removal of which also mostly eliminated Sra1/PIR121 expression (Fig. S1F). Interestingly, other canonical lamellipodial components, including lamellipodin (Lpd) (Dimchev et al., 2020; Law et al., 2013) or the FMNL formin family concluded to operate independently of the Arp2/3 complex machine (Kage et al., 2017) displayed no detectable defects of accumulation at sites of LLS formation (Figs. 4, S5). There was only one exception, the prominent Ena/VASP family of actin polymerases. VASP was the first factor directly associated with promoting actin polymerization at the edge of protrusions such as lamellipodia and at filopodia tips, or the surfaces of bacteria recruiting parts of the actin assembly machinery for intracellular motility (Lambrechts et al., 2008; Laurent et al., 1999; Rottner et al., 1999a; Svitkina et al., 2003). Although changing lamellipodial architecture and size, disruption of all three mammalian Ena/VASP family members did not abolish lamellipodia formation entirely (Damiano-Guercio et al., 2020). Previously observed phenotypes of Ena/VASP loss of function, as for instance the reduction of lamellipodial actin filament mass or width are certainly consistent with the observations described here. Thus, extent, dimension and frequency of formation of LLS described here may well derive from a combination of effects caused by removal of both, WRC-mediated Arp2/3 complex activation and Ena/VASP-dependent promotion of lamellipodial actin assembly. Our data also reveal that all lamellipodial components tested here can be recruited to the cell periphery independent of WRC function, except for Ena/VASP (Figs. 4, S5), which have indeed previously been concluded to operate downstream of WRC, at least in *Dictyostelium* (Litschko et al., 2017). Interestingly, Ena/VASP family member recruitment downstream of WRC might occur through their direct interactions with the WRC subunit Abi established previously (Chen et al., 2014; Litschko et al., 2017). Ena/VASP accumulation in nascent and focal adhesions, however, was unaltered (Fig. S5).

**Figure 3.**
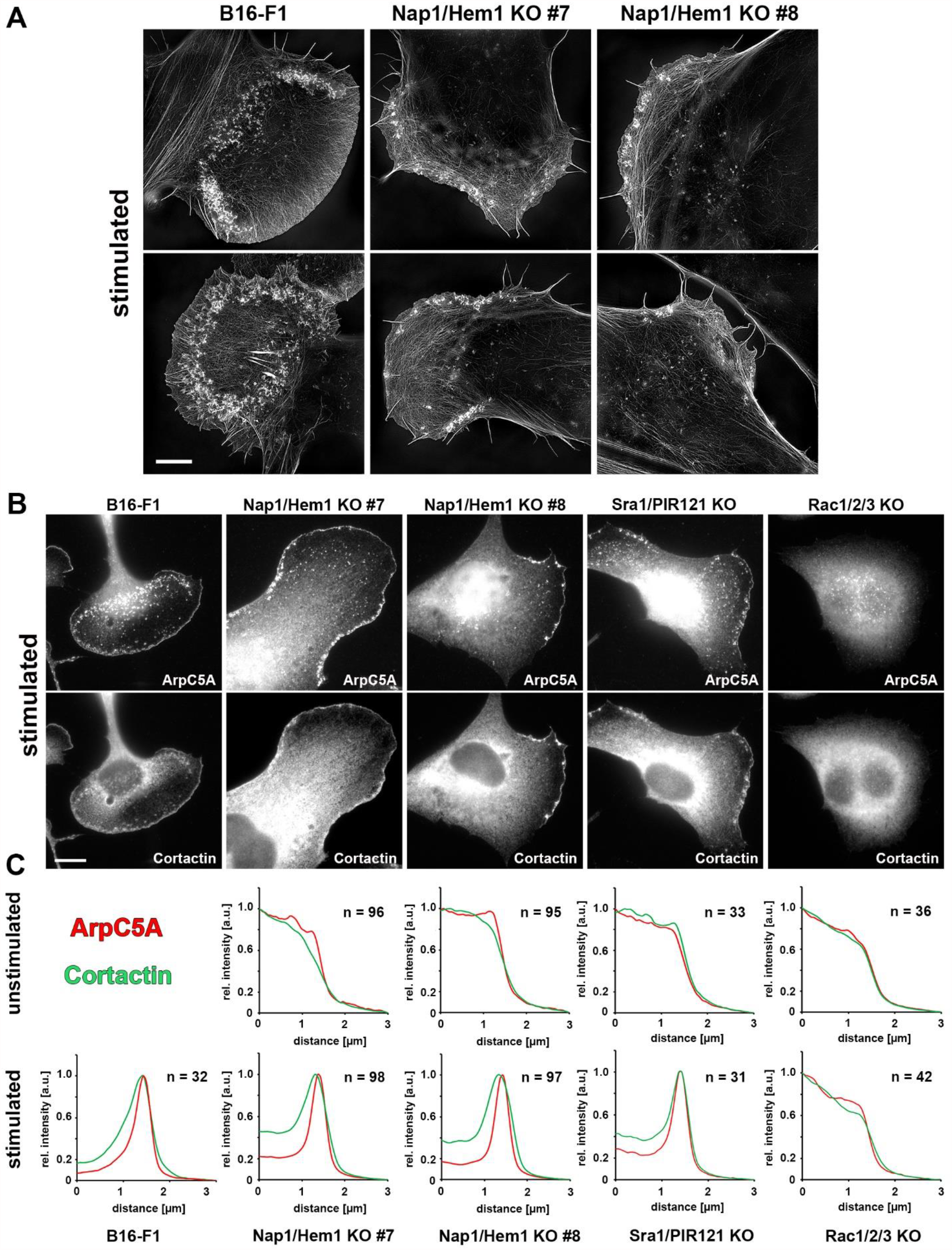
Cells lacking WRC can form lamellipodia-like structures (LLS) upon starvation and stimulation with serum and growth factors. **(A)** Two representative SIM images for each cell line as indicated. Cells were stimulated with HGF in full growth medium, fixed and subjected to phalloidin staining. B16-F1 wildtype cells form characteristically broad, actin-dense lamellipodia with increased dorsal ruffle formation likely effected by stimulation. Scale bar, 10 µm. **(B)** Stimulated cells with genotypes as indicated and stained for endogenous Arp2/3 complex (ArpC5A) and its interactor Cortactin. Note prominent accumulation of respective proteins in lamellipodia of B16-F1 cells and lamellipodia-like structures of WRC-depleted cells, but not in the absence of all three Rac subfamily members (Rac1/2/3 KO cells). Scale bar, 10 µm. **(C)** Graphs show the results from relative line intensity scans averaged from n cells, as indicated. ArpC5A signal intensity (red) and Cortactin signal intensity (green) were measured in cells that were either stimulated or untreated. Measured intensity values were standardized with maximal intensities being normalized to 1 and background signals normalized to 0 for each antibody staining and individual line scan (see Materials and Methods).

**Figure 4.**
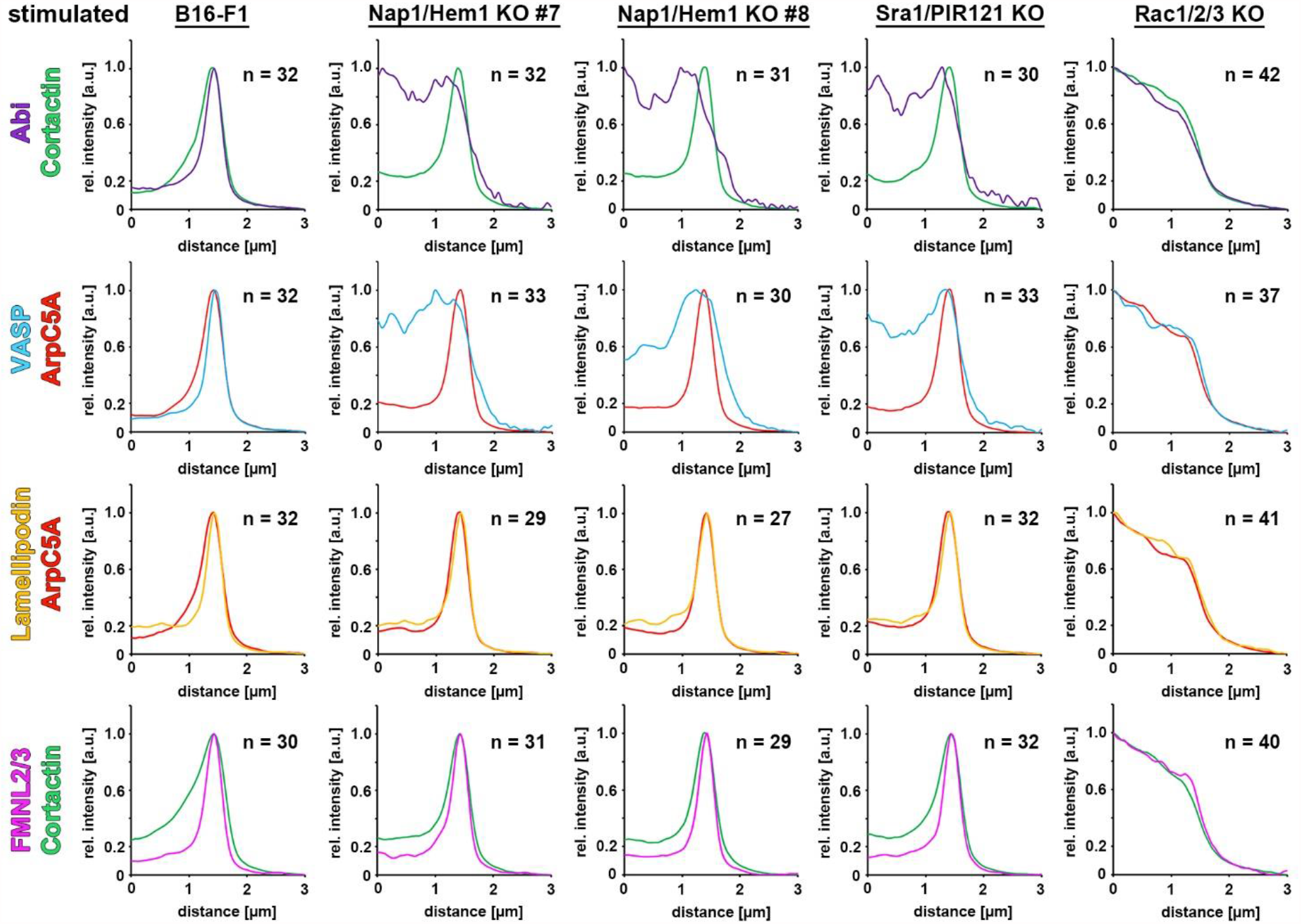
WRC-independent lamellipodia contain all typical lamellipodial tip components except for VASP. Line scan measurements for endogenously-stained lamellipodial tip proteins, such as Abi (purple), VASP (blue), Lamellipodin (orange), FMNL2/3 (pink) in different combinations, and in stimulated cells harboring genotypes as depicted in the figure. Depending on the species of the respective antibody used for counterstainings, either cortactin (polyclonal) or ArpC5A (monoclonal) were used as positive markers for lamellipodia-like structures in WRC-depleted cells. Line scan measurements were performed as described for Fig. 3C. Note the absence of specific VASP accumulation at the tips of lamellipodia in cells of all genotypes except for B16-F1 wildtype. In contrast, none of the components accumulated at the cell peripheries of Rac-deficient B16-F1 (column on the right hand side), showing that LLS are WRC-independent, but Rac-dependent.

### Active Cdc42 can generate cells with striking lamellipodial phenotypes in the absence of WRC

Previous studies have clearly shown contributions of the Rac-related GTPase Cdc42 to protrusion efficiency of and actin assembly in lamellipodia, for instance through FMNL subfamily formins (Kage et al., 2017). The latter pathway, however, was operating independently of Arp2/3 complex-mediated actin filament assembly and branching. In a more recent study, we demonstrated Cdc42 to also be capable of recruiting WRC to the cell periphery in the absence of Rac, at least at low efficiency, but only given that those WRCs did not have to be activated by Rac e.g. due to mutational activation (Schaks et al., 2019). Importantly however, we never observed expression of constitutively active Cdc42 to be capable of stimulating detectable lamellipodia formation in the absence of Rac GTPases, neither in fibroblasts (Steffen et al., 2013) nor in B16-F1 melanoma (Schaks et al., 2019).

However, when expressing constitutively active Cdc42 in WRC-deficient clones of any type used above, we found a subfraction of cells (for quantitations see below) clearly displaying cell morphologies and lamellipodial phenotypes, the molecular composition of which appeared identical to the LLS seen upon serum and growth factor stimulation above (Fig. 5). Aside from actin, these lamellipodial networks again included accumulation of Arp2/3 complex and lamellipodin within or along their edges, but lacked the WRC subunit Abi. As opposed to lamellipodin, VASP was present close to the cell periphery and restricted to numerous nascent, arrow- or dot-shaped adhesions (Fig. 5), but again lacking as continuous line from the lamellipodium edge as common in WT cells (Rottner et al., 1999b; Svitkina et al., 2003).

**Figure 5.**
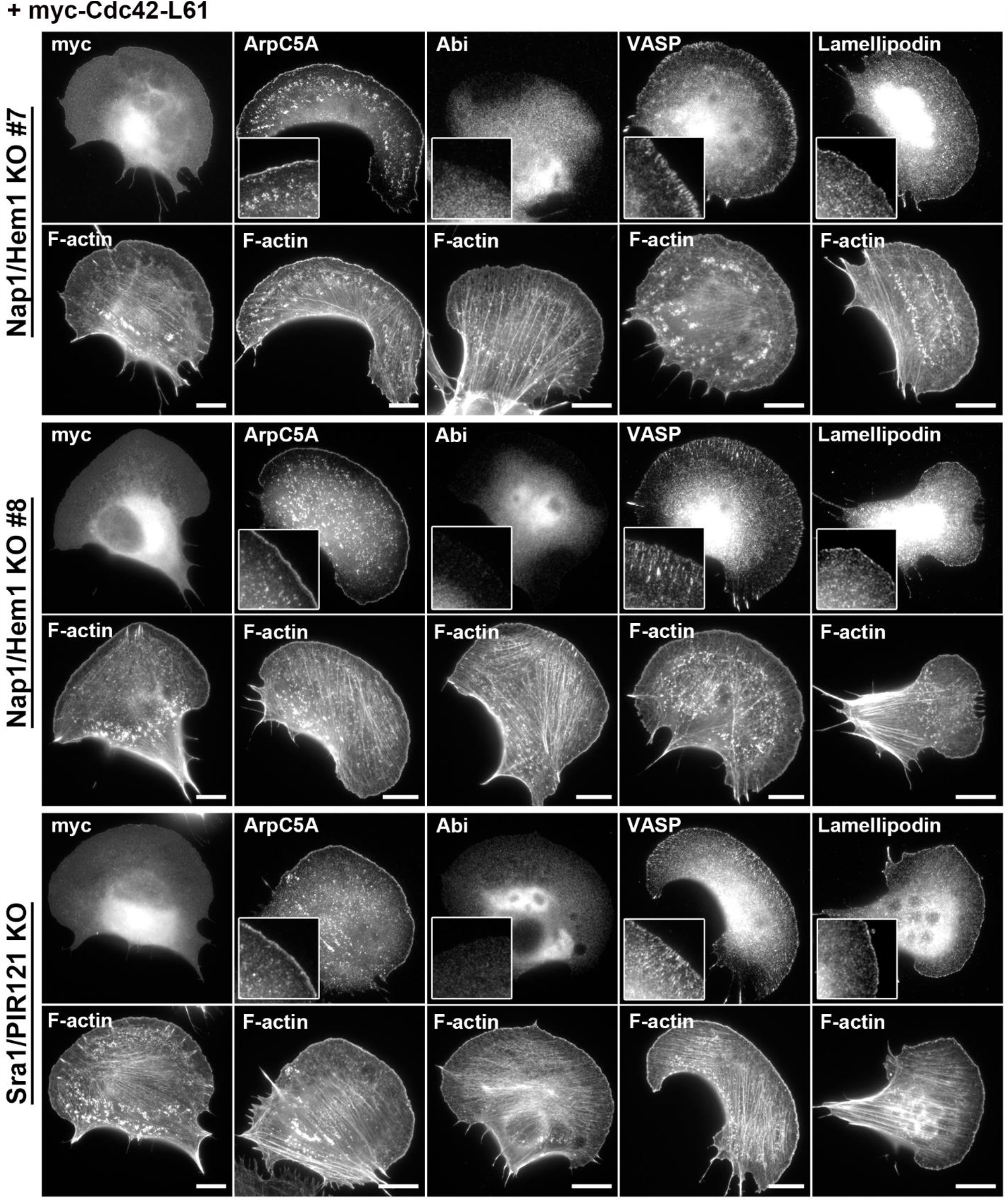
Active Cdc42 can mediate lamellipodia formation despite the lack of WRC. Representative examples of different WRC-depleted cell lines, which were transfected with myc-Cdc42-L61. Expression of the latter was verified by anti-myc staining (left image panel) and overall cell morphology was visualized using phalloidin (F-actin). Note that expression of constitutively-active Cdc42-L61 allows formation of lamellipodia-like structures that are positive for accumulation of canonical markers, such as Arp2/3 complex (ArpC5A) and Lamellipodin (insets), but lack tip enrichment of Abi and VASP (insets). In contrast, VASP localization to nascent and mature focal adhesions is not abrogated by WRC depletion. Scale bar, 10 µm.

### Cdc42-driven lamellipodial phenotypes require its direct effector N-WASP

Our results prompted us to dissect the signaling to Arp2/3 complex-dependent actin assembly in WRC-deficient cells more carefully. To this end, we disrupted either N-WASP or Cdc42 in a B16-F1 clone already lacking functional WRC through removal of Sra1/PIR121 (KO #3, Fig. 6A) (Schaks et al., 2018). We examined two independently generated clones for each novel genotype in more detail (Fig. 6A). Interestingly, side-by-side comparison of cells transfected with myc-tagged, constitutively active Cdc42 revealed that B16-F1 WT and Sra1/PIR121 KO clone #3 displayed similar morphologies of cells positive for the myc-tag (Fig. 6B). However, quantitations revealed that the percentage of cells displaying this phenotype dropped from 100% in WT to approximately 20% in cells lacking WRC (Fig. 6C). This suggested that the majority of lamellipodia formed in B16-F1 WT and additionally boosted by active Cdc42 were likely formed through WRC and concomitant Rac activation, as expected from previous work (Nobes and Hall, 1995; Rottner and Stradal, 2011), but there were clear exceptions not requiring WRC (Fig. 6B, C). Those exceptions formed in Sra1/PIR121-null cells were entirely eliminated upon additional removal of the Cdc42 effector and prominent Arp2/3 complex activator N-WASP (Rohatgi et al., 1999) (Fig. 6B, C). Finally, stimulation of starved cells with the serum/growth factor cocktail used in previous experiments (Figs. 3, 4 and S5) revealed that nearly 12% of WRC-deficient cells displayed LLS, but this population was entirely abolished by additional genetic removal of Cdc42 (Fig. 6D). We conclude that LLS thus constitute peculiar, Arp2/3 complex-containing actin networks formed at the cell periphery, which essentially require both Rac and Cdc42 signaling, and the formation of which is normally superimposed by WRC activity in WT B16-F1 cells. Importantly, these structures can contribute to the efficiency of cell migration, since WRC-deficient cells additionally disrupted for Cdc42 displayed a statistically significant reduction of random cell migration rates (Fig. S6A), which was not seen in B16-F1 clones solely lacking Cdc42 (Figs. S6B, C).

**Figure 6.**
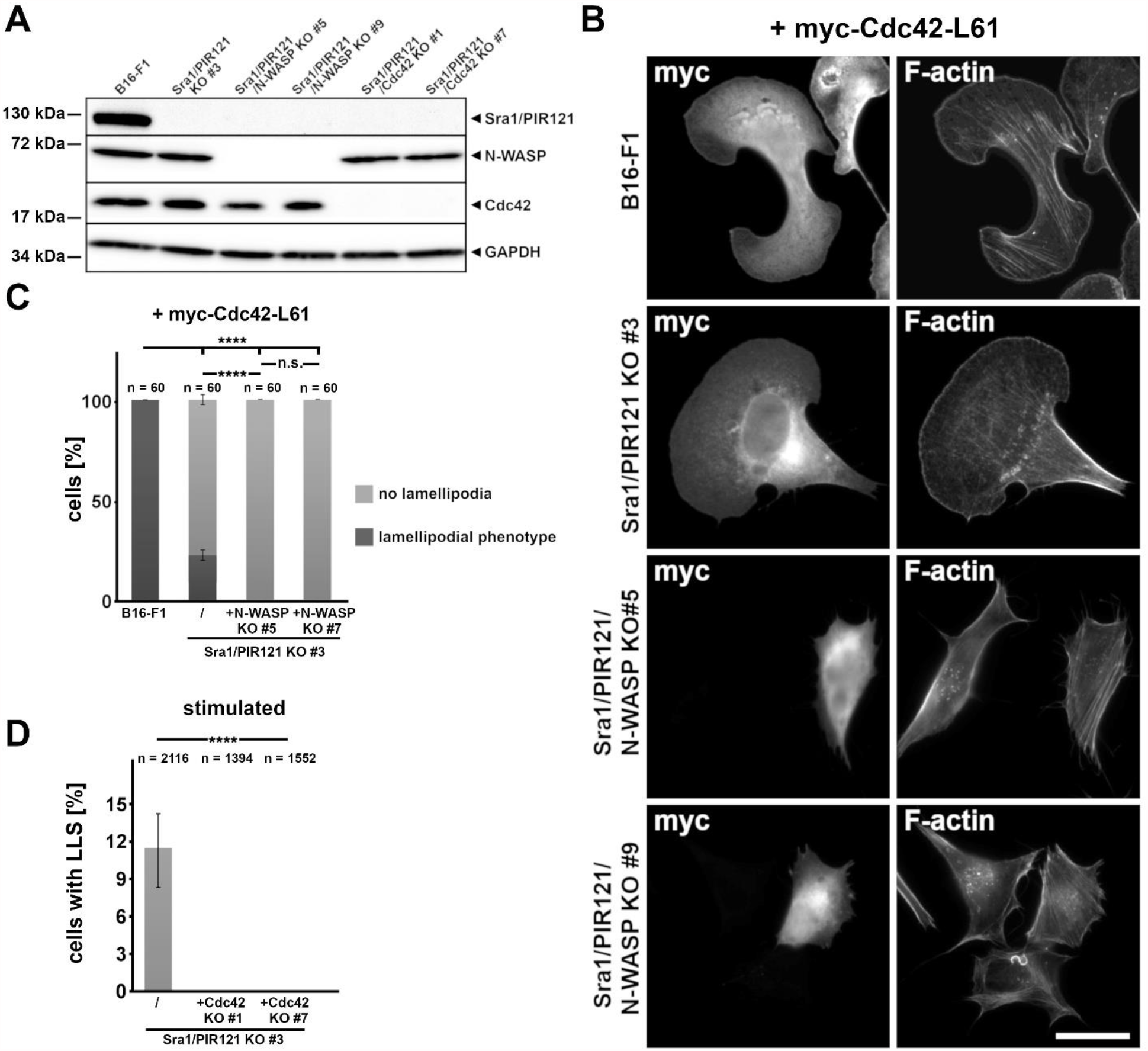
Stimulation of lamellipodia/LLS by active Cdc42 requires N-WASP, and by serum/growth factor treatment Cdc42. **(A)** Western blot of CRISPR/Cas9-modified cell lines employed in this dataset. Removal of Sra1/PIR121 (clone #3; Schaks et al., 2018) was complemented by disruption of N-WASP or Cdc42, as indicated. GAPDH was used as loading control. **(B)** Expression of constitutively active Cdc42 can induce the formation of apparent lamellipodia in both wildtype and WRC-deficient cells (top two panels), but not upon additional removal of N-WASP (2 clones, bottom panels). Cells transfected with myc-tagged, constitutively active Cdc42 (left column) were counterstained for the actin cytoskeleton with phalloidin (right column). Scale bar, 20 µm. **(C)** Quantitation of the data displayed as representative images in B. Note that 100% of B16-F1 wildtype transfected with constitutively active Cdc42 display a lamellipodial phenotype, which is strongly reduced (but not abolished) to app. 20% of cells in the absence of WRC, and abolished in clones additionally lacking N-WASP. **(D)** Quantitation of LLS formation in WRC-deficient cells (Sra1/PIR121 clone #3) upon starvation and stimulation (full medium/HGF), revealing that the LLS formation occurring at a frequency of <12% under these conditions is eliminated upon Cdc42 removal (two independent clones). Statistics in C and D were done using one-way ANOVA with Dunnet’s adjustment for multiple comparisons; ^****^ = p ≤ 0.0001 and n.s. = p ≥ 0.05.

### LLS stimulation and Rac-dependent, but WRC-independent actin remodeling activities are re-capitulated in fibroblasts

In order to examine whether compensatory expression of Hem1 upon Nap1 deletion as well as the peculiar capability of LLS formation in Nap1/Hem1 double KO cells was restricted to B16-F1 melanoma cells, we chose the commonly employed NIH 3T3 fibroblast cell line as independent cell type. In spite of MEFs described to be virtually devoid of lamellipodia upon acute, tamoxifen-mediated deletion of the *Nckap1* gene encoding Nap1 (Whitelaw et al., 2020), isolated NIH 3T3 clones identified to lack Nap1 expression upon CRISPR-treatment by Western blotting (Fig. 7A) still invariably displayed tiny, but clearly detectable lamellipodia (Fig. 7B). Next, we disrupted Hem1 starting from Nap1 KO clone #8, and identified 2 consecutive, Nap1/Hem1 double KO clones clearly devoid of any detectable lamellipodia in regular growth conditions (Fig. 7C). Sequencing of the Hem1 locus confirmed the absence of any wildtype allele (Fig. 7D). Moreover, Western blotting of all five WRC subunits revealed both a clear, but somewhat differential reduction of remaining subunits in the single Nap1 KO #8, and even further reduction if not elimination - like for WAVE - of remaining subunits in both double KO lines (Nap1/Hem1 KOs #2 and #3, Fig. 7E).

**Figure 7.**
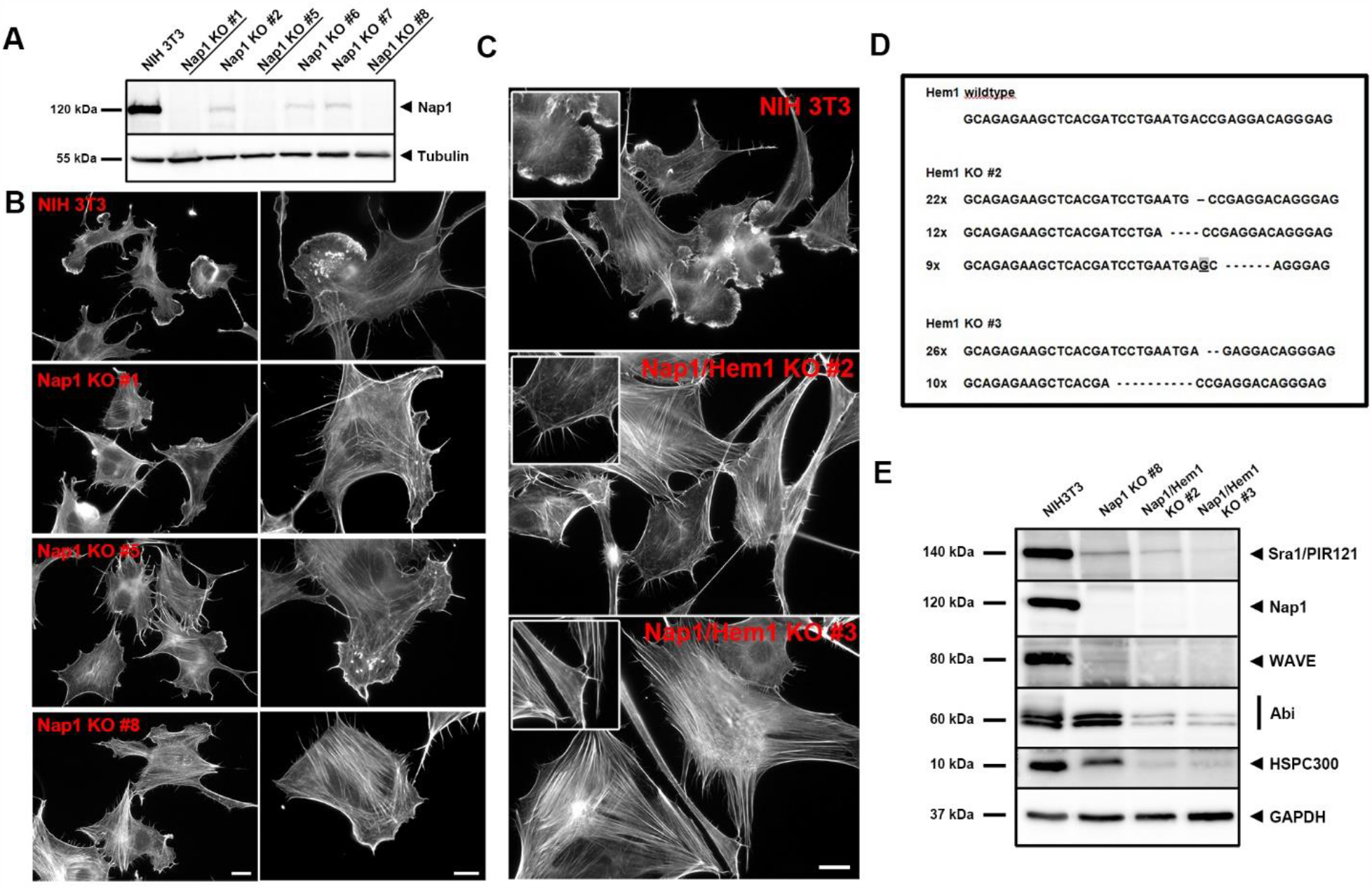
Compensatory Hem1 expression as a consequence of Nap1 KO is independent of cell type. **(A)** Anti-Nap1 western blot confirming CRISPR/Cas9-mediated deletion of Nap1 expression in NIH 3T3 fibroblasts (underlined, clones #1, #5, #8). **(B)** Representative, phalloidin-stained cell images showing Nap1 KO cells forming lamellipodia, although these structures possess compromised morphologies or sizes, at least in part. Overview images, left column (scale bar, 20 µm); higher magnifications, right (scale bar, 10 µm). **(C)** Nap1-deficient clone #8 was subjected to additional, CRISPR-mediated Hem1 removal, causing complete loss of lamellipodia protrusions and increased stress fiber formation in Nap1/Hem double KO cell lines as compared to NIH 3T3 wildtype (insets) in standard growth conditions. Scale bar, 20 µm. **(D)** Sequencing data information from the Hem1 gene locus for the two analyzed clones. Western blot analyses of WRC subunit expression in NIH 3T3 wildtype, Nap1 KO #8 and Nap1/Hem1 double KO clones, as indicated. Note dependence of WRC subunit expression on Nap1/Hem1 gene dose, as illustrated by abrogated WAVE or HSPC expression only in Nap1/Hem1 double KO cell lines.

Strikingly, and as opposed to the phenotypes observed upon regular growth conditions (Fig. 7C), starvation followed by serum plus growth factor stimulation (again HGF in this case) heaved the formation of reasonably prominent LLS in both Nap1/Hem1 double KO clones. Those LLS structures even contained microspike bundles normally restricted to canonical lamellipodia (see insets in Fig. 8A). Assuming that the molecular regulation of these structures would be consistent with our observations already described for B16-F1 melanoma above, we sought instead to explore the physiological relevance of their formation in various complementary assays and model systems for active actin remodeling.

**Figure 8.**
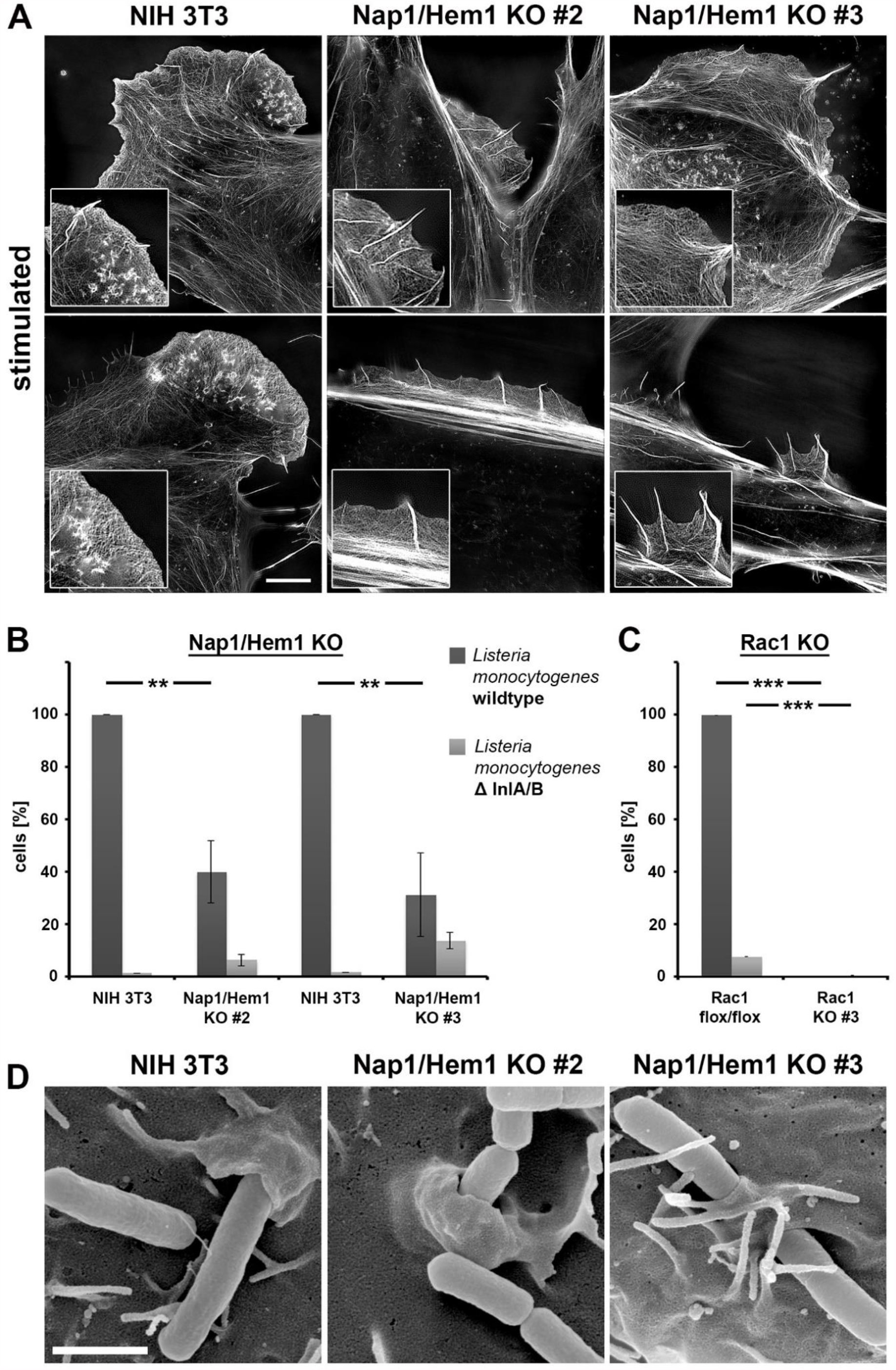
Nap1/Hem1 null fibroblasts are partly responsive to growth factor stimulation and to Listeria-induced, bacterial invasion. **(A)** Structured illumination microscopy images of stimulated cells (HGF/full medium) stained for the actin cytoskeleton with phalloidin (two representative images per cell line). Note that both KO clones, although lacking functional WRC, can form peripheral structures highly reminiscent in appearance and dimension to the lamellipodia in wild type cells (left), demonstrating that this phenomenon is not restricted to B16-F1 cells. **(B)** NIH 3T3 control and KO fibroblasts were tested for invasion efficiency of wildtype *Listeria monocytogenes versus* InternalinA/B‐deficient bacteria (ΔInlA/B) as control, using gentamicin survival assays. Data are arithmetic means ± sem from four independent experiments, each normalized to the uptake of wildtype bacteria respectively obtained with parental NIH 3T3 cells. **(C)** *L*.*m*. invasion assay done as in B, but performed on Rac1 flox/flox or Rac1 KO#3 mouse embryonic fibroblasts lacking any Rac activity (Steffen et al., 2013). Note that loss of WRC reduces *Listeria* invasion to only app. 30-40%, while loss of Rac (Rac1 KO #3) abolishes both specific and non-specific pathogen uptake. Data in B and C were statistically compared using Mann-Whitney rank sum test. ^**^, p ≤ 0.01; ^***^, p ≤ 0.001. **(D)** Representative scanning electron microscopy images showing *L*.*m*. (cylinder-like objects) uptake by cell lines as indicated. In spite of the reduced uptake efficiency upon Nap1/Hem1 removal as compared to control (see B), little changes in the morphology of cell surface structures accompanying bacterial entry could be observed. The variability in the formation frequency of microvilli-like structures in distinct clones was independent of genotype (compare Nap1/Hem1 double KO clones #2 and #3). Scale bar, 1 µm.

The gram-positive bacterium *Listeria monocytogenes* invades host cells making use of direct interaction of bacterial surface or secreted proteins known as internalins. Due to the lack of interaction of Internalin A (InlA) with the murine version of its host cell ligand, E-cadherin, invasion of murine cells is exclusively mediated through Internalin B, which elicits Rac-dependent actin remodeling essential for entry downstream of direct interaction with its ligand on the host cell surface, the HGF receptor also known as c-Met (Pizarro-Cerda et al., 2012). In spite of functions in bacterial entry previously seen for the Rho-GTPase Cdc42, thought to operate in this pathway upstream of Rac signaling, we previously concluded InlB-triggered actin remodeling to require WRC and Nap1 (Bosse et al., 2007). Although we failed to confirm a role in entry for N-WASP in cells genetically deleted for this Cdc42-effector (Bosse et al., 2007), others had previously proposed that elicitation of *Listeria*-induced actin remodeling may involve both WRC and N-WASP (Bierne et al., 2005). Whatever the case, here we clearly demonstrate using our novel Nap1/Hem1-disrupted cell lines that cells lacking functional WRC (Fig. 7E) indeed still can be infected through the InlB/-Met pathway (Fig. 8B), albeit at reduced efficiency as compared to control NIH 3T3 cells. The difference between wildtype *Listeria* and isogenic mutants lacking internalins A and ‐B (ΔInlA/B) proved the specificity of the response for InlB/c-Met signaling. As seen with growth factor-induced formation of LLS in B16-F1 cells, this response was again completely eliminated in fibroblasts lacking Rac GTPases (Steffen et al., 2013), confirming the pattern of Rac-dependent but WRC-independent actin remodeling events uncovered in the context of the current experiments (Fig. 8C). Finally, since the efficiency of bacterial entry was reduced to roughly 30–40% in the absence of WRC, we asked whether those entry events occurring in the absence of WRC were by any means morphologically distinguishable from those in control fibroblasts. However, scanning electron micrographs capturing bacteria right in the middle of the entry process revealed that except for some variability in formation of microvilli-like structures routinely seen in these fibroblast cells and partly accompanying invasion, no major alterations were seen concerning size and shape of the plasma membrane leaflets wrapping around invading bacteria (Fig. 8D). These data are consistent at least with the concept of Listeria invasion being mediated by a so called zipper mechanism of entry (Mostowy and Cossart, 2009), with invading bacteria sinking into their host cells with plasma membrane leaflets zippering around them. It is tempting to speculate that in spite of essential Rac functions, the potency and efficiency of WRC-mediated Arp2/3 complex activation and actin assembly may not be obligatory for these spatially confined actin remodeling events. Notably, these structures are also strongly contrasting the trigger-like plasma membrane projections and ruffles accompanying the invasion of gram-negative bacteria like *Salmonella* and *Shigella* (Rottner et al., 2005).

### WRC-deficient fibroblasts uncover additional actin-remodeling processes differentially requiring WRC *versus* Rac functions

At last, we explored a number of actin dynamics-related phenomena and parameters, which are frequently studied upon interference with actin-binding proteins. Fibroblasts are well known to form circular dorsal ruffles (CDRs), particularly prominently upon growth factor stimulation following cell starvation, and induction of these structures has previously been associated with actin regulators also contributing to lamellipodia formation and peripheral membrane ruffling (Anton et al., 2003; Itoh and Hasegawa, 2013; Lai et al., 2009; Suetsugu et al., 2003). In our previous studies, Rac GTPases turned out to be absolutely obligatory for this process (Steffen et al., 2013), but somewhat surprising, the induction of CDRs appeared even more efficient in Nap1/Hem1 double KO clones as compared to control NIH 3T3 (Fig. S7A, B). To what extent this might be related to potential, compensatory engagement of N-WASP in this process (Legg et al., 2007) will constitute an important aspect of future investigation. In stark contrast, random cell migration efficiency was already markedly reduced upon single Nap1 disruption, and suppressed to about a quarter of the level observed in controls upon Nap1/Hem1 removal (Fig. S7C). These data clearly confirm the relevance of canonical, WRC-dependent lamellipodia for efficient cell migration, as seen already with B16-F1 melanoma (Fig. 2D). Finally, we explored the capability of these cells to chemotax towards a gradient of growth factor, in this case 2.5% serum supplemented with HGF (Fig. S7D). Previous data showed that the formation of Arp2/3 complex-dependent lamellipodia does not constitute an essential prerequisite for efficient chemotaxis (Dimchev et al., 2021; Wu et al., 2012). Considering the absence of at least canonical lamellipodia in Nap1/Hem1 double KO cells confirms this view in an impressive fashion (Fig. S7D), with no reduction in forward migration index in Nap1/Hem1 KO clones (Fig. S7E), and no consistent, genotype-dependent reduction in the rate of chemotactic migration (Fig. S7F). Given the notion that Rac GTPases are absolutely essential for chemotaxis (Steffen et al., 2013), all these data suggest that the relevance of Rac signaling for chemotactic sensing and migration can be clearly separated from its canonical downstream signaling through WRC-mediated Arp2/3 complex activation. Future studies will thus also have to explore how precisely Rac impacts on the fascinating process of chemotaxis.

### Concluding remarks

For nearly two decades now, WRC-dependent, canonical lamellipodia have been the key structures mediating effective cell motility, in particular downstream of Rac GTPase signaling. This was true for an overwhelming majority of studies focusing on canonical migration *in vitro* and in 2D, whereas a seminal study on epithelial tumor cell migration in 3D already uncovered mutual antagonisms between classical, WRC/Arp2/3-dependent migration and N-WASP/Arp2/3-dependent tissue invasion (Tang et al., 2013). Our studies using CRISPR/Cas9-mediated gene editing in various combinations allow us to unravel here that WRC can indeed be bypassed to a certain extent for Arp2/3 complex-dependent actin assembly at the cell periphery even *in vitro* and in 2D, given that specific requirements are fulfilled. However, the WRC-independent structures formed require as essential prerequisite not only the presence of any of the three mammalian Rac GTPases, but also the related GTPase Cdc42. Furthermore, Cdc42 activation by itself can also promote the formation of lamellipodia-like structures in the absence of WRC. The relevance of Cdc42 in this pathway is further emphasized by its gain of impact for cell migration in the absence of WRC. Importantly, our observations are not restricted to migrating melanoma cells, but mirrored in commonly employed fibroblast assays assessing various actin dynamics-dependent processes, including growth factor signaling or stimulated actin remodeling during bacterial entry. Our results highlight once again the importance of choosing the right “factor” if aiming at abolishing specific processes and structures employing genetics.

## Materials and methods

### Cell culture and transfections

B16-F1 murine melanoma cells (ATCC CRL-6323) were cultured in DMEM (4.5 g/L glucose, Invitrogen), supplemented with 10% FCS (Gibco, Paisley, UK) and 2 mM L-glutamine (Thermo Fisher Scientific). Sra1/PIR121 KO and Rac1/2/3 KO cells were as described (Schaks et al., 2018). Transfections of parental B16-F1 cells were carried out in 35 mm dishes using 0.5 µg DNA and 1 µl JetPrime transfection reagent (PolyPlus), whereas 1 µg DNA and 2 µl JetPrime were required to transfect respective KO cell lines. After overnight transfection, B16-F1 cells were seeded onto glass coverslips pre-coated with laminin (25 µg/ml in laminin-coating buffer: 150 mM NaCl, 50 mM Tris, pH 7.4).

NIH 3T3 (ATCC CRL-1658) and derived KO cell lines as well as Rac1^fl/fl^ mouse embryonic fibroblasts and the corresponding Rac1 KO (clone 3) (Steffen et al., 2013) were grown in DMEM (4.5 g/L glucose), 10% FCS (Sigma), 2 mM L-glutamine, 1% non-essential amino acids and 1 mM sodium pyruvate. NIH 3T3 cells and corresponding KO clones were transfected with a total of 3 µg DNA and 9 µl JetPrime in 35 mm dishes overnight. For microscopy, Rac1^fl/fl^ and Rac1 KO MEFs as well as NIH 3T3 cells were seeded in regular growth medium onto coverslips coated with fibronectin (Roche; 25 µg/ml in PBS). All cell lines were cultivated at 37°C in the presence of 7.5% CO_2_.

### Plasmids

EGFP-C1, -C2, -C3 and EGFP-N1, -N2, -N3 vectors used for cloning were purchased from Clontech Inc. (Mountain View, CA, USA). EGFP-tagged, murine Nap1 and -Sra1 were described in (Steffen et al., 2004), and EGFP-N-WASP in (Lommel et al., 2001). EGFP-Hem1 was generated using clone IRAK961M1081Q2 obtained from the German Research Centre for Genome Research (RZPD). PRK5-myc-Cdc42-L61 was a kind gift from Laura Machesky (Beatson Institute, Glasgow, UK). The ΔD21 mutant of EGFP-Hem1 was generated by site-directed mutagenesis using Phusion polymerase employing forward primer 5’-GCTCACGATCCTGAACCGAGGACAGGGAG-3’ and respective complementary sequence as reverse primer.

### CRISPR/Cas9-mediated genome editing

B16-F1 cells as well as NIH 3T3 cells were genome-edited using CRISPR/Cas9 technology in order to disrupt expression of Nap1 (encoded by *NCKAP1*), Hem1 (*NCKAP1L*), N-WASP (*WASL*) or Cdc42 (*CDC42*) in different B16-F1 line backgrounds. Generation of B16-F1 cells eliminated for Nap1 and Sra1/PIR121 expression has already been described in (Dolati et al., 2018) and (Schaks et al., 2018), respectively. Disruption of the *NCKAP1* gene in NIH 3T3 cells was done using a construct harboring the same CRISPR-gRNA sequence as previously described with B16-F1 (5’-GACGCCCCGGTCGTTGAGGA-3’). In this case, isolated and expanded single cell colonies were screened for the absence of Nap1 by Western blotting using polyclonal, Nap1-specific antibodies, and clones lacking detectable Nap1 expression further verified by sequencing of the Nap1 gene locus. To obtain cell lines devoid of both Nap1 and Hem1 expression, consecutive targeting of the Hem1 gene was performed in B16-F1 Nap1 KO clone #21 (Dolati et al., 2018) and NIH 3T3 Nap1 KO clone #8 (this study) using guide sequence 5’-CTCACGATCCTGAATGACCG-3’. Due to the lack of a functional Hem1 antibody, Hem1 KO clones were initially screened for a complete loss of lamellipodia in regular growth conditions as judged upon fixation and staining of the actin cytoskeleton with phalloidin, followed by validation of the loss of alleles giving rise to functional Hem1 expression.

Cdc42-deficient B16-F1 clones were generated using CRISPR-gRNA 5’-ACAATTAAGTGTGTTGTTGT-3’. Sra1/PIR121-deficient B16-F1 clones additionally lacking Cdc42 or N-WASP were generated by targeting Sra1/PIR121 KO clone #3 (Schaks et al., 2018) with aforementioned, Cdc42 CRISPR-gRNA or with CRISPR-gRNA 5’-CACGTTGGTGACCCTCCGCG-3’ (for N-WASP). Screening for clones lacking Cdc42 or N-WASP expression was performed by Western blotting using Cdc42-specific or N-WASP-specific polyclonal antibodies, respectively. Selected clones were validated by sequencing as described above.

Genes were routinely disrupted targeting exon 1, and selected guide sequences cloned into pSpCas9(BB)-2A-Puro (Addgene plasmid ID: 48139). Following overnight transfections (described above), cells were subjected to selection pressure using puromycin (2.5 µg/ml for B16-F1 and 3 µg/ml for NIH 3T3), and then cloned by the limited dilution method (seeding density into 96 well-plates of 0.5 cells/hole).

### CRISPR/Cas9-mediated EGFP knock-in

Nap1 KO subclone #21-12 was chosen for EGFP insertion into the endogenous Hem1 locus, as this was found to exert the highest compensatory Hem1 expression among all clones. CRISPR-guide sequence 5’-TTTGGGTCTTGTTATCCGGA-3’, which is localized shortly upstream of the Hem1 start codon was used for defined targeting of Cas9. The sequence encoding for EGFP as well as app. 300 bp of arms of homology on both sides were ordered to be synthesized by Eurofins Genomics (Ebersberg, Germany). The respective CRISPR-gRNA and linearized insert were co-transfected into cells at a ratio of 1:3, and cells further processed as described in the previous section.

### Western blotting

For preparation of whole cell extracts, cells were washed thrice with ice-cold PBS, lysed using 4x SDS-sample-buffer (Laemmli), boiled for 5 min at 95°C and sonicated to shear genomic DNA. Western blotting was carried out following standard procedures and aside from those mentioned above, using the following, primary antibodies: GAPDH (Calbiochem; clone 6C5; #CB1001; 1:10,000 dilution), Cdc42 (Cell Signaling Technology, #2462, 1:1000 dilution), α-Tubulin (Synaptic Systems, clone 3A2; #302211; 1:5,000 dilution), Sra1/PIR121 (home-made rabbit polyclonal; raised against peptide 4955B; 1:10,000 dilution; Steffen et al., 2004), Nap1 (home-made rabbit polyclonal; raised against peptide 4953B; 1:5,000 dilution; Steffen et al., 2004), Abi (home-made rabbit polyclonal; E3B1; 1:2,000 dilution; Schaks et al., 2018), pan-WAVE (home-made rabbit polyclonal; pAB5502; 1:1000 dilutionSchaks et al., 2018), HSPC300 (Abcam; ab87449; 1:1,000 dilution), N-WASP (home-made rabbit polyclonal; raised against peptide 385-401; 1:400 dilutionLommel et al., 2001), GFP (Roche; #1814460; 1:2,000 dilution), Hem1 (home-made rabbit polyclonal; kindly provided by Dr. Jan Faix; 149; 1:500 dilution; Stahnke et al., 2021).

HRP-conjugated secondary antibodies were anti-mouse IgG (Dianova; #115-035-062; 1:10,000 dilution) or anti-rabbit IgG (Dianova; #111-035-045; 1:10,000 dilution). Chemiluminescence signals were obtained upon incubation with ECL Prime Western Blotting Detection Reagent (GE Healthcare), and recorded with ECL Chemocam imager (Intas, Germany). Densitometric quantification of protein expression as detected by Western blotting was essentially performed following standard procedures using MetaMorph (Molecular Devices, USA). In brief, digitalized background signals were subtracted from specific band intensities and resulting intensities subsequently normalized to those of corresponding loading controls (GAPDH or α-Tubulin, as indicated). Data were plotted as bar charts using Excel 2010 (Microsoft).

### Immunoprecipitation

Commercially available GFP-TrapA (ChromoTek) was used to immunoprecipitate EGFP-tagged Sra1 or EGFP alone as control and probed for potential co-immunoprecipitation of other WRC subunits. B16-F1 cells were transfected overnight as described above. Cells were then lysed using IP-buffer (140 mM KCl, 50 mM Tris-HCl (pH 7.4), 50 mM NaF, 10 mM Na_4_P_2_O_7_, 2 mM MgCl_2_) supplemented with 1% Triton-X100 and a mini complete protease inhibitor pill (Roche). Lysates were centrifuged at 20,000 × g for 15 min at 4 °C. 5 μl of each cell lysate was mixed with SDS sample buffer referred to as input. 30 μl bead slurry was washed 3x and spun down (2,500 × g for 2 min at 4 °C). Subsequently, cell lysates were added to the beads and incubated under constant mixing for 1 h at 4 °C. After centrifugation, 5 μl of each sample was incubated with sample buffer and used as control for unbound, EGFP-tagged proteins (supernatant). Finally, beads were washed thrice with IP-buffer prior to addition of 25 μl 8 × SDS sample buffer. All samples were boiled for 5 min at 95 °C prior to loading onto respective SDS-gels. After blotting proteins onto PVDF membranes (Immobilon), respective antibodies (see ‘Western blotting’) were used to confirm co-immunoprecipitation of endogenous WRC subunits with overexpressed, EGFP-tagged Sra1.

### Immunofluorescence staining

For immunolabeling of proteins of interest, B16-F1 or NIH 3T3 cells were seeded onto glass coverslips coated with laminin (Sigma) or fibronectin (Roche), respectively. Cells were allowed to adhere overnight. On the following day, cells were fixed with pre-warmed, 4% paraformaldehyde (PFA) in PBS for 20 min and permeabilized with 0.05% Triton-X100 (for B16-F1 cells) or 0.1% Triton-X100 (for NIH 3T3 cells) in PBS for 1 min. Prior to antibody staining, cells were blocked with 5% horse serum in 1% BSA/PBS for ∼30 min.

For sole visualization of the actin cytoskeleton using phalloidin, cells were fixed with 0.25% glutaraldehyde in 4% PFA/PBS followed by staining with Alexa Fluor 488-conjugated phalloidin.

Primary antibodies were diluted in 1% BSA/PBS and incubated for 1 h. Used antibodies were as follows: ArpC5A antibody (clone 323H3; undiluted hybridoma supernatant; Olazabal et al., 2002), FMNL2/3-reactive antibody (Abcam; ab57963; 1:20 dilution), Sra1/PIR121 (#2240; 1:20 dilution), Abi (clone W8.3; kindly provided by Dr. Giorgio Scita, Milano, Italy; undiluted hybridoma supernatant), Nap1 (#2391-C; 1:3 dilution; Steffen et al., 2004), HSPC300 (Abcam; ab87449; 1:100 dilution), pan-WAVE (pAB5502; 1:100 dilution; Schaks et al., 2018), Cortactin (Abcam; ab11065-50; 1:100 dilution), Lamellipodin (kindly provided by Dr. Matthias Krause, London, UK; 1:200 dilution; Krause et al., 2004), VASP (5500-A ; 1:300; Jenzora et al., 2005), myc (SIGMA-Aldrich; M5546; clone 9E10; ascites fluid; 1:100 dilution). Primary antibodies were visualized with Alexa Fluor 488- (1:400 dilution) or Alexa Fluor 594-coupled (1:200 dilution) anti-mouse IgG (Invitrogen; #A11029 or #A11032, respectively). Secondary antibodies against rabbit primary antibodies were Alexa Fluor 488- (1:400 dilution) or Alexa Fluor 594-coupled (1:200 dilution) anti-rabbit IgG (Invitrogen, #A11034 or #A11037, respectively). Alexa Fluor 488- and Alexa Fluor 594-coupled phalloidin were obtained from Invitrogen (#A12379 and #A12381, respectively).

### Cell stimulation

For stimulation with serum and growth factors of cells growing on coated coverslips, washing (3x) with PBS was followed by overnight starvation with DMEM lacking any supplements. Next day, cells were treated with prewarmed, regular growth medium for 20 min supplemented with Hepatocyte Growth Factor (HGF), Platelet Derived Growth Factor (PDGF) or Epidermal Growth Factor (EGF), applied in concentrations as follows: HGF (Sigma): 100 ng/ml; PDGF (Sigma): 20 ng/ml; EGF (Sigma): 100 ng/ml. Afterwards, cells were fixed and subjected to immunofluorescence staining as described above.

### Random migration and chemotaxis assay

For random migration assays, cells were seeded subconfluently into laminin/fibronectin-coated µ-slide 4-well phase contrast-optimized, glass bottom microscopy chambers (Ibidi, Martinsried, Germany) using regular growth medium. For chemotaxis assays, µ-Slide Chemotaxis2D chambers (Ibidi, Martinsried, Germany) were employed according to manufacturer’s instructions with adapted cell concentrations and without coating since technically impossible. For cell starvation, NIH 3T3 and derived KOs were subconfluently seeded into the chamber using DMEM without supplements and incubated for ∼6 h. Afterwards, the upper chamber was filled with 2.5% FCS and 100 ng/ml HGF in DMEM (C100) as chemoattractant. The lower chamber was filled with DMEM. Ibidi chambers were mounted onto an inverted microscope (for details see next paragraph), equipped with a 37°C incubator and CO_2_ atmosphere as well as an automated stage. Phase-contrast movies were acquired on multiple, randomly chosen positions using a 10x/0.15 NA Plan Neofluar objective and a frame rate of 12 frames per hour for at least 10 h. For migration speed analyses, cells were manually tracked using ImageJ (https://imagej.nih.gov/ij/), and for determining directionality and mean square displacement of cells, DiPer software was used (Gorelik and Gautreau, 2014).

### Fluorescence recovery after photobleaching

FRAP experiments were performed essentially as described (Dimchev and Rottner, 2018). In brief, cells were mounted in an open chamber (Warner Instruments) and maintained at 37°C with a heater controller (Model TC-324 B, SN 1176) and appropriate pH using Ham’s F12 medium containing all growth supplements. The Axio Observer was equipped with a DG4 light source (Sutter Instrument) for epifluorescence illumination, a VIS-LED for phase-contrast optics, 63x/1.4-NA or 100x/1.4-NA plan apochromatic objectives for high-mag microscopy, and a back-illuminated, cooled, charge-coupled-device (CCD) camera (CoolSnap HQ2, Photometrics) driven by VisiView software (Visitron Systems) for image acquisition. Bleaching was performed using a 405 nm diode laser at 60–70 mW output power) controlled by the 2D-VisiFRAP Realtime Scanner (Visitron Systems, Germany). Comparison of the turnover of EGFP-tagged Nap1 *versus* -Hem1 was performed in the Nap1 KO clone virtually devoid of lamellipodia formation in standard conditions (clone #16). Movies were acquired at app. 2 s per frame (0,5 Hertz). Ectopically expressed, EGFP-tagged proteins were bleached within a lamellipodial region, as indicated in Fig. S3G (red outline). Average intensities in bleached regions were measured over time and background intensities obtained from extracellular regions subtracted for each frame. Acquisition photobleaching was eliminated by normalizing data from each time point to non-bleached, lamellipodial tip regions (green outline in Fig. S3G). Data were curve-fitted with SigmaPlot 12.0 (Scientific Solutions SA, Switzerland) using single exponential equation: f = y0+a*(1-exp(-b*x)), and half times of recovery derived from fitted data.

### Line scan analyses

Fluorescence intensities of the lamellipodial components ArpC5A, Cortactin, VASP, Abi, Lamellipodin and FMNL2/3 (Figures 3 and 4) were measured by drawing an orthogonal line (3 µm long, 5 pxl wide) across the cell edge. Line scan graphs represent corresponding average intensities ranging from the cell center (left) to the outside (right). Line scan measurements were performed using MetaMorph software (Molecular Devices, USA) and further processed using Microsoft Excel. At least 30 cells for each cell clone and distinct staining were analyzed and averaged. Background intensities outside the cell were subtracted, values were normalized according to the maximal/minimal intensity measured for each condition and plotted over the entire line distance of 3 µm.

### 3D – Structured Illumination Microscopy

Images were acquired on a Nikon SIM-E superresolution microscope equipped with a LU-N3-SIM 488/561/640 laser unit mounted on a Nikon Ti eclipse (Nikon) and a Piezo z drive (Mad city labs). Images were taken using a CFI Apochromat TIRF 100x/1.49 NA oil immersion objective (Nikon), a Hamamatsu Orca flash 4.0 LT camera and an N-SIM motorized quad band filter combined with N-SIM 488 and 561 bandpass emission filters using laser line 488 driven by NIS-Elements software (Nikon). Reconstructions were performed employing the slice reconstruction tool (Nikon, NIS-Elements).

### Listeria invasion assay

Bacterial invasion with *Listeria monocytogenes* (EGD wt) and isogenic, specific invasion-deficient control bacteria (ΔInlAB) (Parida et al., 1998) into wildtype NIH and corresponding Nap1/Hem1 double KO clones or Rac1^flox/flox^ and corresponding Rac1 KO#3 fibroblasts (Steffen et al., 2013) was quantified by gentamycin protection assay essentially as described previously (Bosse et al., 2007). The only exception was that as opposed to previous protocols (Elsinghorst, 1994), host cell stress was minimized by performing infections in the presence of 2% serum as opposed to DMEM alone. Data from three and four independent experiments for Rac KO and Nap1/Hem1 KO clones, respectively, were displayed as normalized to 100% of wildtype host cell control in each case, and subjected to statistical analyses as specified below.

### Scanning electron microscopy

Field emission scanning electron microscopy: For the visualization of invasion events with *Listeria monocytogenes*, we used a modified version of a previously described assay (Rohde et al., 2003). In brief, NIH control (5 x 10^4^ cells) or Nap1/Hem1 double KO clones (1 x 10^5^ cells) were seeded onto 12 mm coverslips in 24-well plate holes and allowed to grow for 18-20 hours before infections. Infections were performed with *L. monocytogenes* diluted 1:100 into DMEM/2% FBS after overnight culture in BHI-medium and a gentle washing step in DMEM. Invasion was stopped at various time points after start of infections by adding fixative to the growth medium (final concentration: 5% formaldehyde and 2.5% glutaraldehyde) in order to maximize the probability of capturing active entry events. After fixation, coverslips were washed twice with TE buffer (20 mM Tris-HCl, 1 mM EDTA, pH 6.9) and dehydrated in 10 minute-steps at increased acetone concentrations on ice (10, 30, 50, 70, 90%), followed by two steps at 100% and room temperature. Cells were critical-point dried with liquid CO_2_(CPD 30, Balzers, Liechtenstein) and covered with a palladium–gold film by sputter coating (SCD 500, Bal-Tec, Liechtenstein) before examination in a field emission scanning electron microscope (Zeiss Merlin, Oberkochen, Germany) using the HESE2 Everhart Thornley SE detector and the in-lens SE detector in a 25:75 ratio at an acceleration voltage of 5 kV.

### Data Processing and Statistical Analyses

Brightness and contrast levels of images were adjusted using MetaMorph software. Figures were further processed and assembled with Photoshop CS4. Data analyses were carried out in ImageJ and MetaMorph, Excel 2010 and Sigma plot 12.0. Statistical comparisons of multiple datasets were done using one-way ANOVA with post-hoc Tukey (Prism 6.01). In case of pair-wise comparisons, non-parametric Mann-Whitney rank sum test was used (Sigma plot 12.0). A probability of error of 5% (p ≤ 0.05; * in Figure panels) was considered to indicate statistical significance. **, *** and **** indicated p-values ≤ 0.01, 0.001 and 0.0001, respectively.

## Acknowledgments

We thank Drs. Jan Faix, Matthias Krause and Giorgio Scita for kindly providing reagents. This work was supported in part by the Deutsche Forschungsgemeinschaft (DFG), Research Training Group GRK2223 and individual grants RO2414/3-2 (to K.R.) and KA5106/1-1 (to F.K.). We would like to thank Brigitte Denker and Ina Schleicher for excellent technical assistance.

## Author contributions

Conceptualization and Methodology, K.R., F.K., A.S., M.S., and M.Mü.; Investigations, F.K., H.D., M.Mi., M.S., F.G., S.S., and M.Mü.; Visualization, F.K. and H.D.; Writing – Original Draft, K.R. and F.K.; Writing – Review & Editing, K.R., F.K. and T.E.B.S.; Funding Acquisition, K.R. and T.E.B.S., Supervision, K.R.

## Declaration of Interests

The authors declare no competing interests.

## Notes

### Competing Interest Statement

The authors have declared no competing interest.

## References

Anton, I.M., S.P. Saville, M.J. Byrne, C. Curcio, N. Ramesh, J.H. Hartwig, and R.S. Geha. 2003. WIP participates in actin reorganization and ruffle formation induced by PDGF. J Cell Sci. 116:2443–2451.

Aspenstrom, P., A. Fransson, and J. Saras. 2004. Rho GTPases have diverse effects on the organization of the actin filament system. Biochem J. 377:327–337.

Ballestrem, C., B. Wehrle-Haller, and B.A. Imhof. 1998. Actin dynamics in living mammalian cells. J Cell Sci. 111 (Pt 12):1649–1658.

Bear, J.E., J.F. Rawls, and C.L. Saxe, 3rd. 1998. SCAR, a WASP-related protein, isolated as a suppressor of receptor defects in late Dictyostelium development. J Cell Biol. 142:1325–1335.

Bierne, H., H. Miki, M. Innocenti, G. Scita, F.B. Gertler, T. Takenawa, and P. Cossart. 2005. WASP-related proteins, Abi1 and Ena/VASP are required for Listeria invasion induced by the Met receptor. J Cell Sci. 118:1537–1547.

Bosse, T., J. Ehinger, A. Czuchra, S. Benesch, A. Steffen, X. Wu, K. Schloen, H.H. Niemann, G. Scita, T.E. Stradal, C. Brakebusch, and K. Rottner. 2007. Cdc42 and phosphoinositide 3-kinase drive Rac-mediated actin polymerization downstream of c-Met in distinct and common pathways. Mol Cell Biol. 27:6615–6628.

Chen, B., H.T. Chou, C.A. Brautigam, W. Xing, S. Yang, L. Henry, L.K. Doolittle, T. Walz, and M.K. Rosen. 2017. Rac1 GTPase activates the WAVE regulatory complex through two distinct binding sites. Elife. 6.

Chen, X.J., A.J. Squarr, R. Stephan, B. Chen, T.E. Higgins, D.J. Barry, M.C. Martin, M.K. Rosen, S. Bogdan, and M. Way. 2014. Ena/VASP proteins cooperate with the WAVE complex to regulate the actin cytoskeleton. Dev Cell. 30:569–584.

Damiano-Guercio, J., L. Kurzawa, J. Mueller, G. Dimchev, M. Schaks, M. Nemethova, T. Pokrant, S. Bruhmann, J. Linkner, L. Blanchoin, M. Sixt, K. Rottner, and J. Faix. 2020. Loss of Ena/VASP interferes with lamellipodium architecture, motility and integrin-dependent adhesion. Elife. 9.

Davidson, A.J., and R.H. Insall. 2013. SCAR/WAVE: A complex issue. Commun Integr Biol. 6:e27033.

Dimchev, G., B. Amiri, A.C. Humphries, M. Schaks, V. Dimchev, T.E.B. Stradal, J. Faix, M. Krause, M. Way, M. Falcke, and K. Rottner. 2020. Lamellipodin tunes cell migration by stabilizing protrusions and promoting adhesion formation. J Cell Sci. 133.

Dimchev, G., and K. Rottner. 2018. Micromanipulation Techniques Allowing Analysis of Morphogenetic Dynamics and Turnover of Cytoskeletal Regulators. J Vis Exp.

Dimchev, V., I. Lahmann, S.A. Koestler, F. Kage, G. Dimchev, A. Steffen, T.E.B. Stradal, F. Vauti, H.H. Arnold, and K. Rottner. 2021. Induced Arp2/3 Complex Depletion Increases FMNL2/3 Formin Expression and Filopodia Formation. Front Cell Dev Biol. 9:634708.

Dolati, S., F. Kage, J. Mueller, M. Musken, M. Kirchner, G. Dittmar, M. Sixt, K. Rottner, and M. Falcke. 2018. On the relation between filament density, force generation, and protrusion rate in mesenchymal cell motility. Mol Biol Cell. 29:2674–2686.

Eden, S., R. Rohatgi, A.V. Podtelejnikov, M. Mann, and M.W. Kirschner. 2002. Mechanism of regulation of WAVE1-induced actin nucleation by Rac1 and Nck. Nature. 418:790-793.

Elsinghorst, E.A. 1994. Measurement of invasion by gentamicin resistance. Methods Enzymol. 236:405–420.

Gallop, J.L. 2020. Filopodia and their links with membrane traffic and cell adhesion. Semin Cell Dev Biol. 102:81–89.

Gorelik, R., and A. Gautreau. 2014. Quantitative and unbiased analysis of directional persistence in cell migration. Nat Protoc. 9:1931–1943.

Innocenti, M., A. Zucconi, A. Disanza, E. Frittoli, L.B. Areces, A. Steffen, T.E. Stradal, P.P. Di Fiore, M.F. Carlier, and G. Scita. 2004. Abi1 is essential for the formation and activation of a WAVE2 signalling complex. Nat Cell Biol. 6:319–327.

Itoh, T., and J. Hasegawa. 2013. Mechanistic insights into the regulation of circular dorsal ruffle formation. J Biochem. 153:21–29.

Jenzora, A., B. Behrendt, J.V. Small, J. Wehland, and T.E. Stradal. 2005. PREL1 provides a link from Ras signalling to the actin cytoskeleton via Ena/VASP proteins. FEBS Lett. 579:455–463.

Kage, F., M. Winterhoff, V. Dimchev, J. Mueller, T. Thalheim, A. Freise, S. Bruhmann, J. Kollasser, J. Block, G. Dimchev, M. Geyer, H.J. Schnittler, C. Brakebusch, T.E. Stradal, M.F. Carlier, M. Sixt, J. Kas, J. Faix, and K. Rottner. 2017. FMNL formins boost lamellipodial force generation. Nat Commun. 8:14832.

Kobayashi, K., S. Kuroda, M. Fukata, T. Nakamura, T. Nagase, N. Nomura, Y. Matsuura, N. Yoshida-Kubomura, A. Iwamatsu, and K. Kaibuchi. 1998. p140Sra-1 (specifically Rac1-associated protein) is a novel specific target for Rac1 small GTPase. J Biol Chem. 273:291–295.

Krause, M., and A. Gautreau. 2014. Steering cell migration: lamellipodium dynamics and the regulation of directional persistence. Nat Rev Mol Cell Biol. 15:577–590.

Krause, M., J.D. Leslie, M. Stewart, E.M. Lafuente, F. Valderrama, R. Jagannathan, G.A. Strasser, D.A. Rubinson, H. Liu, M. Way, M.B. Yaffe, V.A. Boussiotis, and F.B. Gertler. 2004. Lamellipodin, an Ena/VASP ligand, is implicated in the regulation of lamellipodial dynamics. Dev Cell. 7:571–583.

Kunda, P., G. Craig, V. Dominguez, and B. Baum. 2003. Abi, Sra1, and Kette control the stability and localization of SCAR/WAVE to regulate the formation of actin-based protrusions. Curr Biol. 13:1867–1875.

Lai, F.P., M. Szczodrak, J.M. Oelkers, M. Ladwein, F. Acconcia, S. Benesch, S. Auinger, J. Faix, J.V. Small, S. Polo, T.E. Stradal, and K. Rottner. 2009. Cortactin promotes migration and platelet-derived growth factor-induced actin reorganization by signaling to Rho-GTPases. Mol Biol Cell. 20:3209–3223.

Lambrechts, A., K. Gevaert, P. Cossart, J. Vandekerckhove, and M. Van Troys. 2008. Listeria comet tails: the actin-based motility machinery at work. Trends Cell Biol. 18:220–227.

Laurent, V., T.P. Loisel, B. Harbeck, A. Wehman, L. Grobe, B.M. Jockusch, J. Wehland, F.B. Gertler, and M.F. Carlier. 1999. Role of proteins of the Ena/VASP family in actin-based motility of Listeria monocytogenes. J Cell Biol. 144:1245–1258.

Law, A.L., A. Vehlow, M. Kotini, L. Dodgson, D. Soong, E. Theveneau, C. Bodo, E. Taylor, C. Navarro, U. Perera, M. Michael, G.A. Dunn, D. Bennett, R. Mayor, and M. Krause. 2013. Lamellipodin and the Scar/WAVE complex cooperate to promote cell migration in vivo. J Cell Biol. 203:673–689.

Legg, J.A., G. Bompard, J. Dawson, H.L. Morris, N. Andrew, L. Cooper, S.A. Johnston, G. Tramountanis, and L.M. Machesky. 2007. N-WASP involvement in dorsal ruffle formation in mouse embryonic fibroblasts. Mol Biol Cell. 18:678–687.

Leithner, A., A. Eichner, J. Muller, A. Reversat, M. Brown, J. Schwarz, J. Merrin, D.J. de Gorter, F. Schur, J. Bayerl, I. de Vries, S. Wieser, R. Hauschild, F.P. Lai, M. Moser, D. Kerjaschki, K. Rottner, J.V. Small, T.E. Stradal, and M. Sixt. 2016. Diversified actin protrusions promote environmental exploration but are dispensable for locomotion of leukocytes. Nat Cell Biol. 18:1253–1259.

Litschko, C., J. Linkner, S. Bruhmann, T.E.B. Stradal, T. Reinl, L. Jansch, K. Rottner, and J. Faix. 2017. Differential functions of WAVE regulatory complex subunits in the regulation of actin-driven processes. Eur J Cell Biol. 96:715–727.

Lommel, S., S. Benesch, K. Rottner, T. Franz, J. Wehland, and R. Kuhn. 2001. Actin pedestal formation by enteropathogenic Escherichia coli and intracellular motility of Shigella flexneri are abolished in N-WASP-defective cells. EMBO Rep. 2:850–857.

Machesky, L.M., and R.H. Insall. 1998. Scar1 and the related Wiskott-Aldrich syndrome protein, WASP, regulate the actin cytoskeleton through the Arp2/3 complex. Curr Biol. 8:1347–1356.

McCarty, O.J., M.K. Larson, J.M. Auger, N. Kalia, B.T. Atkinson, A.C. Pearce, S. Ruf, R.B. Henderson, V.L. Tybulewicz, L.M. Machesky, and S.P. Watson. 2005. Rac1 is essential for platelet lamellipodia formation and aggregate stability under flow. J Biol Chem. 280:39474–39484.

Miki, H., H. Yamaguchi, S. Suetsugu, and T. Takenawa. 2000. IRSp53 is an essential intermediate between Rac and WAVE in the regulation of membrane ruffling. Nature. 408:732–735.

Mostowy, S., and P. Cossart. 2009. Cytoskeleton rearrangements during Listeria infection: clathrin and septins as new players in the game. Cell Motil Cytoskeleton. 66:816–823.

Nakagawa, H., H. Miki, M. Nozumi, T. Takenawa, S. Miyamoto, J. Wehland, and J.V. Small. 2003. IRSp53 is colocalised with WAVE2 at the tips of protruding lamellipodia and filopodia independently of Mena. J Cell Sci. 116:2577–2583.

Nobes, C.D., and A. Hall. 1995. Rho, rac, and cdc42 GTPases regulate the assembly of multimolecular focal complexes associated with actin stress fibers, lamellipodia, and filopodia. Cell. 81:53–62.

Olazabal, I.M., E. Caron, R.C. May, K. Schilling, D.A. Knecht, and L.M. Machesky. 2002. Rho-kinase and myosin-II control phagocytic cup formation during CR, but not FcgammaR, phagocytosis. Curr Biol. 12:1413–1418.

Parida, S.K., E. Domann, M. Rohde, S. Muller, A. Darji, T. Hain, J. Wehland, and T. Chakraborty. 1998. Internalin B is essential for adhesion and mediates the invasion of Listeria monocytogenes into human endothelial cells. Mol Microbiol. 28:81–93.

Park, H., M.M. Chan, and B.M. Iritani. 2010. Hem-1: putting the “WAVE” into actin polymerization during an immune response. FEBS Lett. 584:4923–4932.

Pizarro-Cerda, J., A. Kuhbacher, and P. Cossart. 2012. Entry of Listeria monocytogenes in mammalian epithelial cells: an updated view. Cold Spring Harb Perspect Med. 2.

Rogers, S.L., U. Wiedemann, N. Stuurman, and R.D. Vale. 2003. Molecular requirements for actin-based lamella formation in Drosophila S2 cells. J Cell Biol. 162:1079–1088.

Rohatgi, R., L. Ma, H. Miki, M. Lopez, T. Kirchhausen, T. Takenawa, and M.W. Kirschner. 1999. The interaction between N-WASP and the Arp2/3 complex links Cdc42-dependent signals to actin assembly. Cell. 97:221–231.

Rohde, M., E. Muller, G.S. Chhatwal, and S.R. Talay. 2003. Host cell caveolae act as an entry-port for group A streptococci. Cell Microbiol. 5:323–342.

Rottner, K., B. Behrendt, J.V. Small, and J. Wehland. 1999a. VASP dynamics during lamellipodia protrusion. Nat Cell Biol. 1:321–322.

Rottner, K., J. Faix, S. Bogdan, S. Linder, and E. Kerkhoff. 2017. Actin assembly mechanisms at a glance. J Cell Sci. 130:3427–3435.

Rottner, K., A. Hall, and J.V. Small. 1999b. Interplay between Rac and Rho in the control of substrate contact dynamics. Curr Biol. 9:640–648.

Rottner, K., and M. Schaks. 2019. Assembling actin filaments for protrusion. Curr Opin Cell Biol. 56:53–63.

Rottner, K., and T.E. Stradal. 2011. Actin dynamics and turnover in cell motility. Curr Opin Cell Biol. 23:569–578.

Rottner, K., T.E. Stradal, and J. Wehland. 2005. Bacteria-host-cell interactions at the plasma membrane: stories on actin cytoskeleton subversion. Dev Cell. 9:3–17.

Rotty, J.D., C. Wu, and J.E. Bear. 2013. New insights into the regulation and cellular functions of the ARP2/3 complex. Nat Rev Mol Cell Biol. 14:7–12.

Salzer, E., S. Zoghi, M.G. Kiss, F. Kage, C. Rashkova, S. Stahnke, M. Haimel, R. Platzer, M. Caldera, R.C. Ardy, B. Hoeger, J. Block, D. Medgyesi, C. Sin, S. Shahkarami, R. Kain, V. Ziaee, P. Hammerl, C. Bock, J. Menche, L. Dupre, J.B. Huppa, M. Sixt, A. Lomakin, K. Rottner, C.J. Binder, T.E.B. Stradal, N. Rezaei, and K. Boztug. 2020. The cytoskeletal regulator HEM1 governs B cell development and prevents autoimmunity. Sci Immunol. 5.

Schaks, M., H. Doring, F. Kage, A. Steffen, T. Klunemann, W. Blankenfeldt, T. Stradal, and K. Rottner. 2019. RhoG and Cdc42 can contribute to Rac-dependent lamellipodia formation through WAVE regulatory complex-binding. Small GTPases:1–11.

Schaks, M., S.P. Singh, F. Kage, P. Thomason, T. Klunemann, A. Steffen, W. Blankenfeldt, T.E. Stradal, R.H. Insall, and K. Rottner. 2018. Distinct Interaction Sites of Rac GTPase with WAVE Regulatory Complex Have Non-redundant Functions in Vivo. Curr Biol. 28:3674–3684 e3676.

Snapper, S.B., F. Takeshima, I. Anton, C.H. Liu, S.M. Thomas, D. Nguyen, D. Dudley, H. Fraser, D. Purich, M. Lopez-Ilasaca, C. Klein, L. Davidson, R. Bronson, R.C. Mulligan, F. Southwick, R. Geha, M.B. Goldberg, F.S. Rosen, J.H. Hartwig, and F.W. Alt. 2001. N-WASP deficiency reveals distinct pathways for cell surface projections and microbial actin-based motility. Nat Cell Biol. 3:897–904.

Stahnke, S., H. Doring, C. Kusch, D.J.J. de Gorter, S. Dutting, A. Guledani, I. Pleines, M. Schnoor, M. Sixt, R. Geffers, M. Rohde, M. Musken, F. Kage, A. Steffen, J. Faix, B. Nieswandt, K. Rottner, and T.E.B. Stradal. 2021. Loss of Hem1 disrupts macrophage function and impacts migration, phagocytosis, and integrin-mediated adhesion. Curr Biol. 31:2051–2064 e2058.

Steffen, A., J. Faix, G.P. Resch, J. Linkner, J. Wehland, J.V. Small, K. Rottner, and T.E. Stradal. 2006. Filopodia formation in the absence of functional WAVE-and Arp2/3-complexes. Mol Biol Cell. 17:2581–2591.

Steffen, A., S.A. Koestler, and K. Rottner. 2014. Requirements for and consequences of Rac-dependent protrusion. Eur J Cell Biol. 93:184–193.

Steffen, A., M. Ladwein, G.A. Dimchev, A. Hein, L. Schwenkmezger, S. Arens, K.I. Ladwein, J. Margit Holleboom, F. Schur, J. Victor Small, J. Schwarz, R. Gerhard, J. Faix, T.E. Stradal, C. Brakebusch, and K. Rottner. 2013. Rac function is crucial for cell migration but is not required for spreading and focal adhesion formation. J Cell Sci. 126:4572–4588.

Steffen, A., K. Rottner, J. Ehinger, M. Innocenti, G. Scita, J. Wehland, and T.E. Stradal. 2004. Sra-1 and Nap1 link Rac to actin assembly driving lamellipodia formation. EMBO J. 23:749–759.

Stradal, T.E.B., and M. Schelhaas. 2018. Actin dynamics in host-pathogen interaction. FEBS Lett. 592:3658–3669.

Suetsugu, S., D. Yamazaki, S. Kurisu, and T. Takenawa. 2003. Differential roles of WAVE1 and WAVE2 in dorsal and peripheral ruffle formation for fibroblast cell migration. Dev Cell. 5:595–609.

Svitkina, T.M., E.A. Bulanova, O.Y. Chaga, D.M. Vignjevic, S. Kojima, J.M. Vasiliev, and G.G. Borisy. 2003. Mechanism of filopodia initiation by reorganization of a dendritic network. J Cell Biol. 160:409–421.

Tang, H., A. Li, J. Bi, D.M. Veltman, T. Zech, H.J. Spence, X. Yu, P. Timpson, R.H. Insall, M.C. Frame, and L.M. Machesky. 2013. Loss of Scar/WAVE complex promotes N-WASP-and FAK-dependent invasion. Curr Biol. 23:107–117.

Tang, Q., M. Schaks, N. Koundinya, C. Yang, L.W. Pollard, T.M. Svitkina, K. Rottner, and B.L. Goode. 2020. WAVE1 and WAVE2 have distinct and overlapping roles in controlling actin assembly at the leading edge. Mol Biol Cell. 31:2168–2178.

Veltman, D.M., J.S. King, L.M. Machesky, and R.H. Insall. 2012. SCAR knockouts in Dictyostelium: WASP assumes SCAR’s position and upstream regulators in pseudopods. J Cell Biol. 198:501–508.

Whitelaw, J.A., K. Swaminathan, F. Kage, and L.M. Machesky. 2020. The WAVE Regulatory Complex Is Required to Balance Protrusion and Adhesion in Migration. Cells. 9.

Wu, C., S.B. Asokan, M.E. Berginski, E.M. Haynes, N.E. Sharpless, J.D. Griffith, S.M. Gomez, and J.E. Bear. 2012. Arp2/3 is critical for lamellipodia and response to extracellular matrix cues but is dispensable for chemotaxis. Cell. 148:973–987.

Zhu, Z., Y. Chai, Y. Jiang, W. Li, H. Hu, W. Li, J.W. Wu, Z.X. Wang, S. Huang, and G. Ou. 2016. Functional Coordination of WAVE and WASP in C. elegans Neuroblast Migration. Dev Cell. 39:224–238.

