## Supplementary figures and legends for "Lamellipodia-like actin networks in cells lacking WAVE Regulatory Complex"

[1]Division of Molecular Cell Biology, Zoological Institute, Technische Universität Braunschweig, Spielmannstrasse 7, 38106 Braunschweig, Germany; [2]Department of Cell Biology, Helmholtz Centre for Infection Research, Inhoffenstrasse 7, 38124 Braunschweig, Germany; [3]Central Facility for Microscopy, Helmholtz Center for Infection Research HZI, Braunschweig; Braunschweig Integrated Centre of Systems Biology (BRICS), Braunschweig, Germany; [5]present address: Department of Biochemistry and Cell Biology, Geisel School of Medicine at Dartmouth, Hanover, NH, USA

#### Legends to Supplementary Figures

**Figure S1. Nap1 disruption can cause compensatory upregulation of Hem1 expression. (A)** CRISPR/Cas9-mediated loss of Nap1 expression as documented by Western blotting. Two clones (#1 and #2) harbored a faint, residual expression of Nap1 protein, but clones #6, #14, #16, #21 and #23 were completely devoid of detectable Nap1 expression and selected for further analyses (underlined). α-Tubulin was probed for equal sample loading. **(B)** Sequencing results of the CRISPR/Cas9-modified Nap1 gene locus (exon 1) of respective cell clones. Numbers on the left illustrate the frequencies of given mutations found, indicating that all clones aside from clone #21 likely possess the same insertion on all alleles present. Yet, all alleles were confirmed to result in frame-shifts and premature stop-codons (not shown). **(C)** Quantitation of expression levels of WRC subunits in Nap1 KO clones as compared to B16-F1 wildtype cells. Bar charts represent relative protein expression measured on ECL exposed membranes normalized to Tubulin from 4 independently generated lysates for each clone. Error bars are ± standard errors of means (sem). Note that loss of Nap1 is accompanied by significant reduction of Sra1 and WAVE levels, while Abi and HSPC300 levels are less affected (the latter found not to be statistically significant throughout all clones). Statistics were done using one-way ANOVA with Dunnet's adjustment for multiple comparisons; **** = $p \leq 0.0001$, n.s (not significant) = $p \geq 0.05$. **(D)** Co-immunoprecipitation of endogenous WRC subunits including Hem1 with EGFP-tagged Sra1 in Nap1 KO #21 (right panel), but not in B16-F1 cells or in either cell line using EGFP alone as control (left panel). Note that endogenous WRC subunits are also robustly precipitated in B16-F1 wildtype cells (exemplified by WAVE and Abi), but as opposed to Nap1 KO #21 (right panel), those precipitates lacked Hem1, likely due to lack of detectable Hem1 expression in B16-F1 wildtype cells. **(E)** Genomic DNA sequencing of the *hem1* locus upon generation of Nap1/Hem1 double KO cell lines (double KO clones #7 and #8, as indicated), revealing various mutations in the Hem1 gene locus (exon 1). In both clones, three distinct mutations on apparently at least three distinct alleles were found, most of which caused premature stop signals. The in-frame deletion (3 nucleotides) found on one allele present in Nap1/Hem1 KO clone #7 was confirmed to represent a loss-of-function mutation (see Fig. S4). **(F)** Quantitation of expression of remaining WRC subunits in B16-F1 wildtype *versus* Nap1/Hem1 double KO clones #7 and -#8, as derived from 6 independently generated cell extracts. Although suppression of remaining WRC subunit expression was clearly differential, reduction was statistically significant for all subunits; one-way ANOVA with Dunnet's adjustment for multiple comparisons; **** = $p \leq 0.0001$, ** = $p \leq 0.01$.

**Figure S2. Analysis of Nap1-depleted cells by structured illumination microscopy (SIM).** SIM images of different Nap1 KO clones and B16-F1 control cells (two representative images each) stained for the filamentous actin cytoskeleton (F-actin). While almost all wildtype B16-F1 cells form lamellipodia on laminin, subfractions of Nap1 KO clones form lamellipodia (left example for all clones), whereas the majority of cells extend filopodia and harbor actin-rich clusters near cell peripheries. Note that the lamellipodia in Nap1 null clones are commonly thinner and less F-actin-dense, except for those observed in Nap1 KO #21.

**Figure S3. Nap1 and Hem1 have redundant functions in lamellipodia formation and cell migration. (A)** Comparison of lamellipodia formation rescued by transient expression of either EGFP-Nap1 or -Hem1 in two clones virtually devoid of lamellipodia formation (B16-F1 clones #16 and #23). Images display EGFP fluorescence of respective construct (green in merges on the right) and the actin cytoskeleton counterstained with phalloidin (red in merges). Scale bars, 10 µm. **(B)** Quantification of cells harboring lamellipodia upon expression of constructs as in A. Data in the bar chart are arithmetic means ± sem from three independent experiments. Number and morphology of such reconstituted lamellipodia were indistinguishable between Nap1 and Hem1-expressing cells. **(C)** Cell lysates of Nap1- or Hem1-transfected cells compared to EGFP-expressing cells confirmed equal rescue of expression of endogenous WRC subunits, like Sra1 and WAVE, upon expression of EGFP-Nap1 *versus* EGFP-Hem1. α-Tubulin was used as loading control. **(D)** B16-F1 control as well as Nap1 KO cells re-expressing either Nap1 or Hem1 were analyzed for random migration speed. Box and whisker plots represent data as follows: boxes correspond to 50% of data points (25%–75%), and whiskers correspond to 80% (10%–90%). Outliers are shown as dots, red numbers in boxes correspond to medians. Statistics in B and D were done using one-way ANOVA with Dunnet's adjustment for multiple comparisons; **** = $p \leq 0.0001$, * = $p \leq 0.05$ and n.s. = $p \geq 0.05$. **(E)** Graph displaying changes in migration directionality over elapsed time (min) for cell lines as illustrated by the color code. Directionality is displayed as the ratio of displacement to trajectory length for each time point. Error bars are ± sem. **(F)** Mean square displacement ($\mu m^2$) over elapsed time (min) for indicated cells lines. Error bars for each time point are ± sem. Note the marked but highly comparable increase of MSD for cell populations expressing EGFP-tagged Nap1 *versus* -Hem1. **(G)** Time-lapse frames of Nap1- or Hem1-expressing cells subjected to FRAP. Time is given in seconds. First image illustrates individual regions that were used for bleaching of fluorescent protein at the lamellipodium tip (red outline), for determination of acquisition photo-bleaching in a reference region (green outline) and background intensity (cyan circle). Bars are 10 µm in overviews and 3 µm in insets. **(H)** FRAP analysis of either EGFP-Nap1 or -Hem1 expressed in Nap1 KO cells. Raw data (blue curve) are plotted as arithmetic means ± sem of fluorescence intensities at acquired time points after bleaching. Half-times of recovery (red numbers) for each component were derived from respective curve fits (red lines).

**Figure S4. A single Hem1 amino acid depletion disrupts its lamellipodia formation capability. (A)** Nap1/Hem1 CRISPR/Cas9-treated clones were probed for rescue of lamellipodia formation with EGFP-tagged, wildtype Hem1 or an in-frame deletion mutant lacking aspartate at position 21, as endogenously observed among other variants in Nap1/Hem1 double KO #7 (see below). While wildtype Hem1 efficiently rescued lamellipodia formation in both cell lines, as confirmed by F-actin staining using phalloidin (right panels), mutated Hem1ΔD21 completely failed to induce lamellipodia protrusions. Scale bar, 10 µm. **(B)** Comparison of genomic sequences and their translated amino acid residues in the relevant region of exon 1 in wildtype *versus* Nap1/Hem1 KO #7, which is devoid of lamellipodia formation in standard conditions (see also Fig. 1F). Note that the in-frame deletion in Nap1/Hem1 double KO #7 among other alleles causes the loss of aspartate at position 21 (Hem1ΔD21). **(C)** Western blot using EGFP-specific antibodies to confirm expression of the Hem1ΔD21-construct as full length protein at expected molecular weight, excluding that its lack of capability to mediate lamellipodia formation is caused by failure of proper expression. α-Tubulin was used as loading control.

**Figure S5. Lamellipodia-like structures are enriched for canonical lamellipodial tip markers Lamellipodin and FMNL2/3, but not for VASP.** Starved and stimulated (HGF in full medium) cells harboring genotypes as depicted on top of the image. Cells with lamellipodia-like protrusions (LLS) or B16-F1 cells harboring canonical lamellipodia were stained for various lamellipodia markers as indicated on the left. Rac1/2/3 KO cells that are incapable of forming these protrusions are shown to confirm specificity of each antibody, and with the exception of VASP in focal adhesions, display no comparable accumulations at the cell periphery. Lamellipodial markers were co-stained for either ArpC5A (monoclonal) or Cortactin (polyclonal), dependent on species of respective, combined antibody. Note that VASP is also enriched in peripheral focal adhesions, but absent from the tips of LLS in WRC-depleted cells. These stainings are representative to the ones subjected to line scan analyses shown in Fig. 4.

**Figure S6. Cdc42 removal reduces migration of B16-F1 cells in the absence of WRC, but not in its presence. (A)** Random migration assay using combined WRC and Cdc42 KO cells established and characterized in Fig. 6 as compared to their parental, control cell line lacking WRC alone (Sra1/PIR121 KO #3). Both clones additionally eliminated for Cdc42 expression display migration rates that were reduced in a statistically significant fashion. **(B)** Western blotting confirming the loss of Cdc42 expression upon CRISPR/Cas9-mediated targeting of the *Cdc42* locus in the wildtype background (three independent Cdc42 KO clones analyzed, as indicated). **(C)** Random migration efficiency of B16-F1 wildtype *versus* Cdc42 KO cells, confirming that Cdc42 loss of function does not significantly impact on migration rates in the presence of WRC. Detailed characterization of the new Cdc42 KO lines will be described elsewhere. Statistics in A and C were done using one-way ANOVA with Dunnet's adjustment for multiple comparisons; **** = $p \leq 0.0001$, ** = $p \leq 0.01$, n.s (not significant) = $p \geq 0.05$.

**Figure S7. Nap1/Hem1 KO fibroblasts display increased, stimulation-induced dorsal ruffling, abrogated rates of migration, but no defects in maintaining directionality during chemotaxis. (A)** Representative phalloidin images of NIH 3T3 wild type and Nap1/Hem1 KO

#3 upon starvation or stimulation with full medium containing distinct growth factors as indicated (+HGF, EGF or PDGF). Note the robust induction of dorsal ruffles upon stimulation, and in particular in the absence of WRC. Scale bar, 20 µm. **(B)** Quantitation of cells with or without dorsal ruffles and in conditions as described in A, but with both Nap1/Hem1 double KO clones (#2 and #3) as compared to wildtype NIH 3T3. Approximately 500 cells were analyzed for each cell line and treatment. Data were collected from three independent experiments and are expressed as arithmetic mean ± sem. **(C)** Box and whiskers plot representing random migration speed of cell lines as denoted in the image. Data displayed as described for Fig. S3D. **(D)** Chemotactic migration towards a gradient of 2.5% serum and 100 ng/ml HGF, as described previously (Steffen et al., 2013). Data sets are from three independent experiments. Rose plots with 10° segments, displaying the frequency of directions of given migration paths during chemotaxis. **(E)** Forward migration index of NIH 3T3 and Nap1/Hem1 KO cell lines during chemotaxis. **(F)** Box and whiskers plots showing medians of chemotactic migration rates in given cell lines. In spite of variability in migration efficiency among Nap1/Hem1 double KO clones, no genotype-specific trend in chemotactic migration efficiency with or without WRC could be discerned. Statistics in B, C, E, F were done using one-way ANOVA with Dunnet's adjustment for multiple comparisons; **** = $p \leq 0.0001$, *** = $p \leq 0.001$, ** = $p \leq 0.01$, * = $p \leq 0.05$, n.s = $p \geq 0.05$.

#### Supplementary Reference

Steffen, A., M. Ladwein, G.A. Dimchev, A. Hein, L. Schwenkmezger, S. Arens, K.I. Ladwein, J. Margit Holleboom, F. Schur, J. Victor Small, J. Schwarz, R. Gerhard, J. Faix, T.E. Stradal, C. Brakebusch, and K. Rottner. 2013. Rac function is crucial for cell migration but is not required for spreading and focal adhesion formation. *J Cell Sci*. 126:4572-4588.

### Figure S1

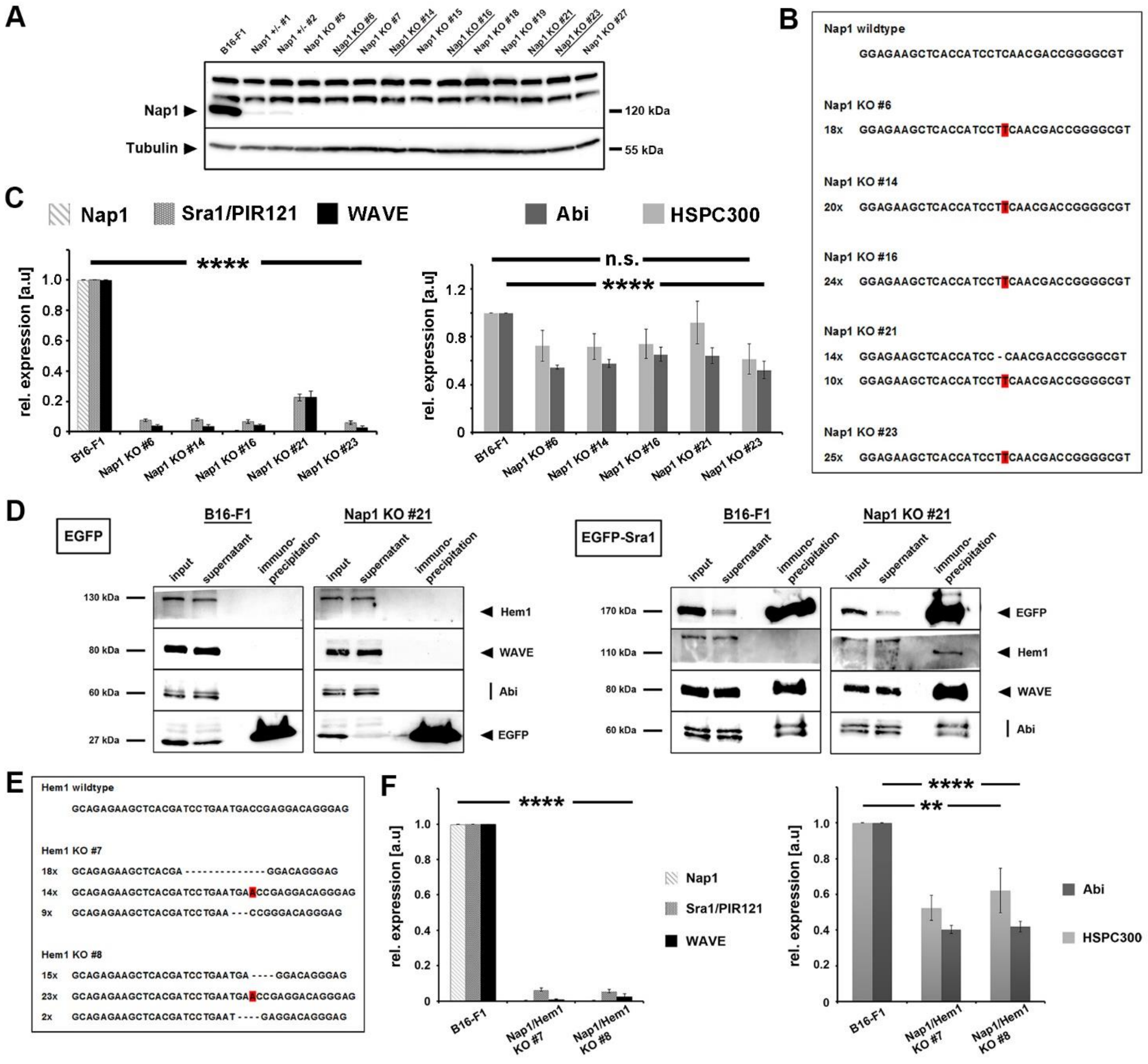
A
B16-F1
Nap1 +/- #1
Nap1 +/- #2
Nap1 KO #5
Nap1 KO #6
Nap1 KO #7
Nap1 KO #14
Nap1 KO #15
Nap1 KO #16
Nap1 KO #18
Nap1 KO #19
Nap1 KO #21
Nap1 KO #23
Nap1 KO #27
Nap1
Tubulin
120 kDa
55 kDa
B
Nap1 wildtype
GGAGAAGCTCACCATCCTCAACGACCGGGGCGT
Nap1 KO #6
18x GGAGAAGCTCACCATCCTTCAACGACCGGGGCGT
Nap1 KO #14
20x GGAGAAGCTCACCATCCTTCAACGACCGGGGCGT
Nap1 KO #16
24x GGAGAAGCTCACCATCCTTCAACGACCGGGGCGT
Nap1 KO #21
14x GGAGAAGCTCACCATCC - CAACGACCGGGGCGT
10x GGAGAAGCTCACCATCCTTCAACGACCGGGGCGT
Nap1 KO #23
25x GGAGAAGCTCACCATCCTTCAACGACCGGGGCGT
C
Nap1
Sra1/PIR121
WAVE
Abi
HSPC300
rel. expression [a.u]
****
n.s.
B16-F1
Nap1 KO #6
Nap1 KO #14
Nap1 KO #16
Nap1 KO #21
Nap1 KO #23
D
EGFP
EGFP-Sra1
B16-F1
Nap1 KO #21
input
supernatant
immuno-precipitation
130 kDa
80 kDa
60 kDa
27 kDa
170 kDa
110 kDa
Hem1
WAVE
Abi
EGFP
E
Hem1 wildtype
GCAGAGAAGCTCACGATCCTGAATGACCGAGGACAGGGAG
Hem1 KO #7
18x GCAGAGAAGCTCACGA - - - - - - - - - - - - - - GGACAGGGAG
14x GCAGAGAAGCTCACGATCCTGAATGAACCGAGGACAGGGAG
9x GCAGAGAAGCTCACGATCCTGAA - - - CCGGGACAGGGAG
Hem1 KO #8
15x GCAGAGAAGCTCACGATCCTGAATGA - - - - GGACAGGGAG
23x GCAGAGAAGCTCACGATCCTGAATGAACCGAGGACAGGGAG
2x GCAGAGAAGCTCACGATCCTGAAT - - - - GAGGACAGGGAG
F
****
**
Nap1/Hem1 KO #7
Nap1/Hem1 KO #8

**Figure S2**

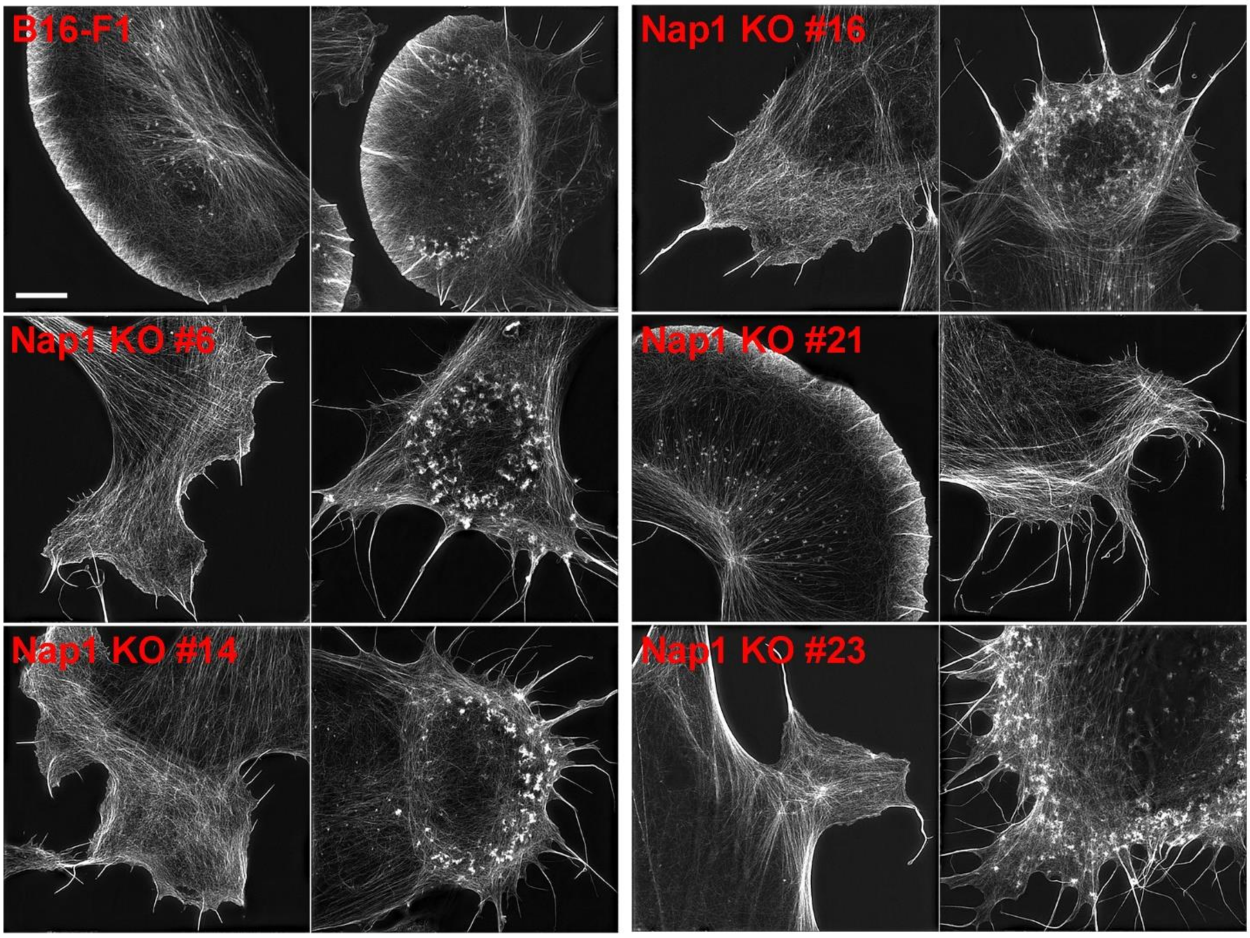

#### Figure S3

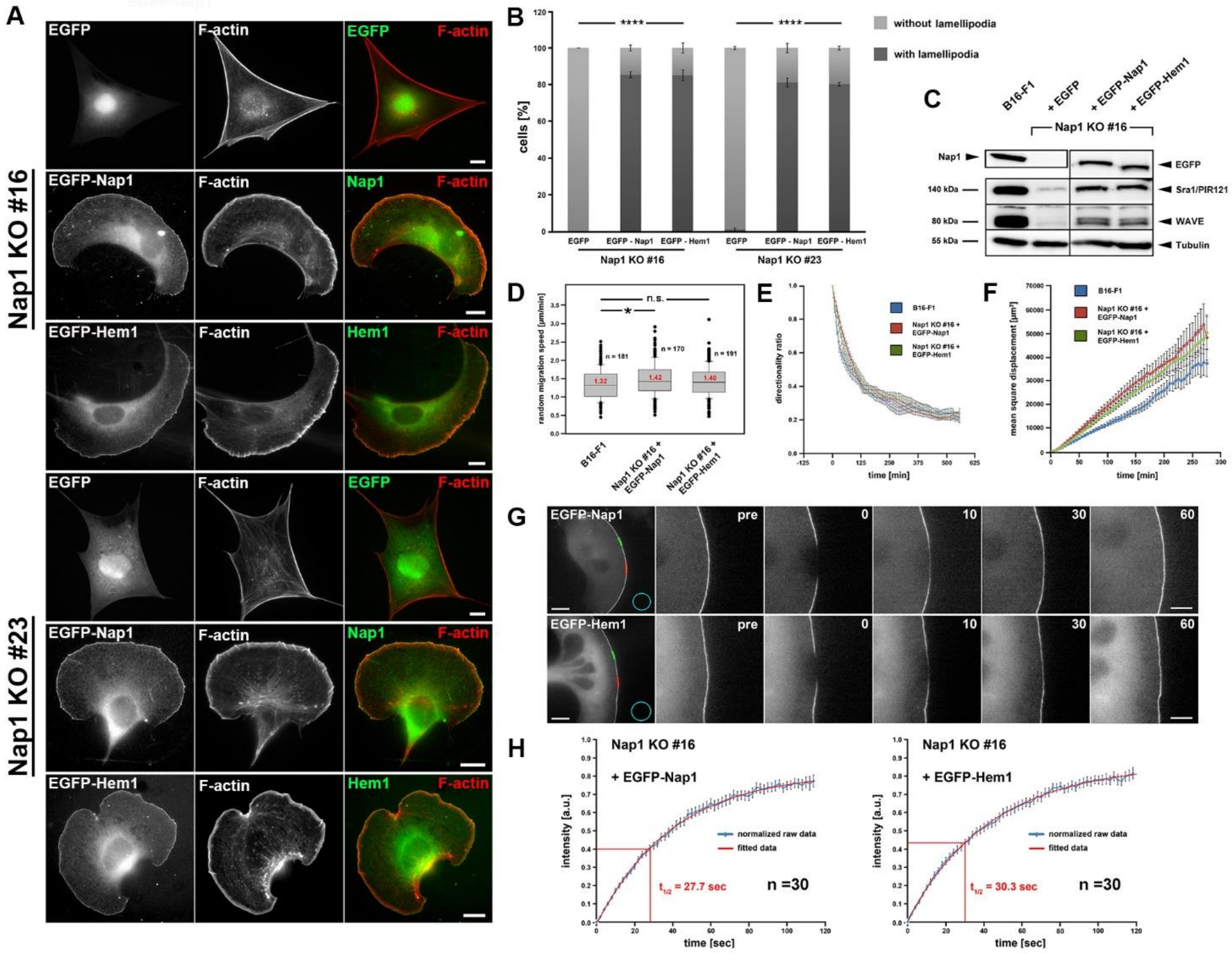
A
EGFP
F-actin
EGFP F-actin
Nap1 KO #16
EGFP-Nap1
Nap1 F-actin
EGFP-Hem1
Hem1 F-actin
Nap1 KO #23
B
cells [%]
without lamellipodia
with lamellipodia
EGFP
EGFP - Nap1
EGFP - Hem1
Nap1 KO #16
Nap1 KO #23
C
B16-F1
+ EGFP
+ EGFP-Nap1
+ EGFP-Hem1
Nap1 KO #16
Nap1
140 kDa
80 kDa
55 kDa
EGFP
Sra1/PIR121
WAVE
Tubulin
D
random migration speed [µm/min]
n.s.
n = 181
n = 170
n = 191
1.32
1.42
1.40
B16-F1
Nap1 KO #16 + EGFP-Nap1
Nap1 KO #16 + EGFP-Hem1
E
directionality ratio
time [min]
F
mean square displacement [µm²]
G
pre
0
10
30
60
H
Nap1 KO #16
+ EGFP-Nap1
+ EGFP-Hem1
intensity [a.u.]
normalized raw data
fitted data
t1/2 = 27.7 sec
t1/2 = 30.3 sec
n =30
time [sec]

**Figure S4**

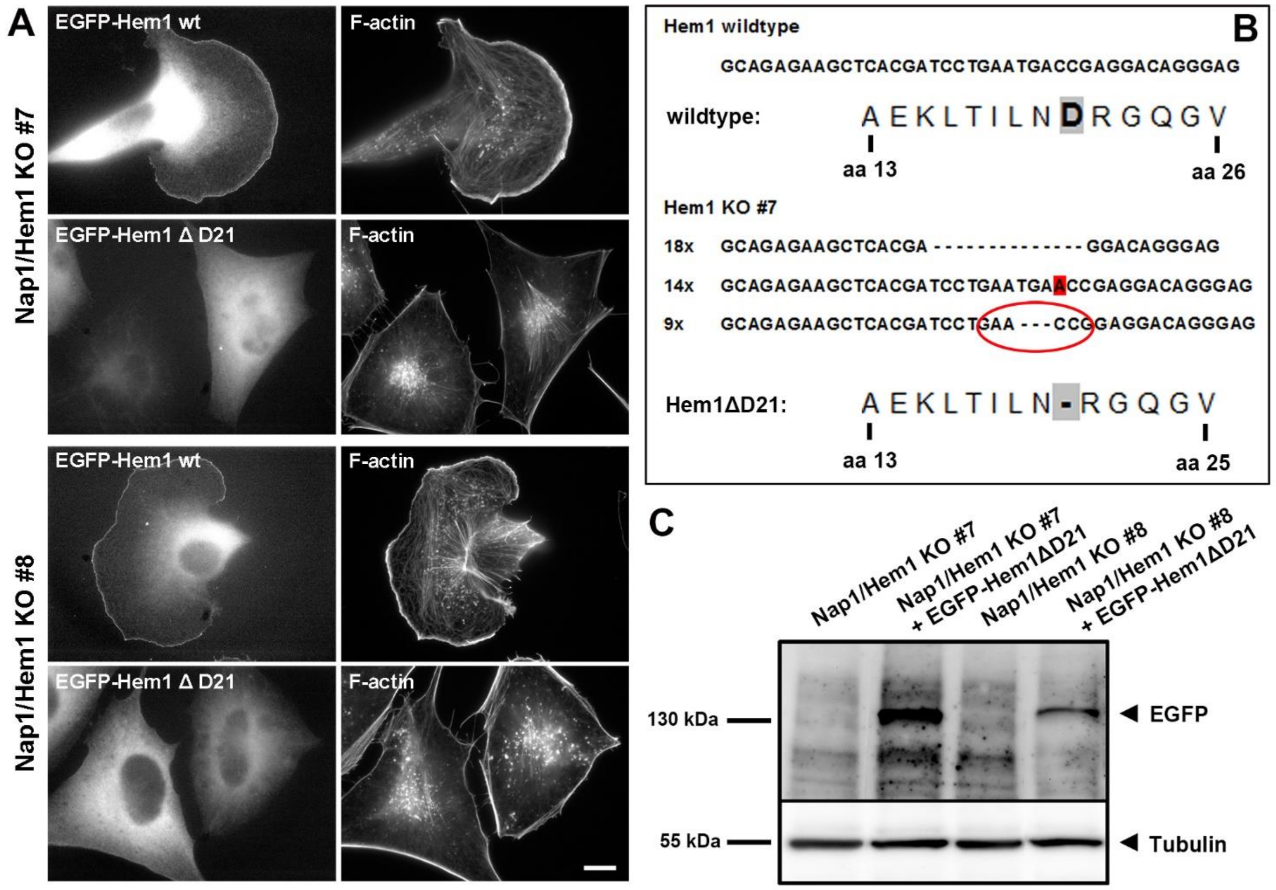

A
EGFP-Hem1 wt
F-actin
EGFP-Hem1 Δ D21
Nap1/Hem1 KO #7
Nap1/Hem1 KO #8
B
Hem1 wildtype
GCAGAGAAGCTCACGATCCTGAATGACCGAGGACAGGGAG
wildtype:
A E K L T I L N D R G Q G V
aa 13
aa 26
Hem1 KO #7
18x GCAGAGAAGCTCACGA - - - - - - - - - - - - - - GGACAGGGAG
14x GCAGAGAAGCTCACGATCCTGAATGAACCGAGGACAGGGAG
9x GCAGAGAAGCTCACGATCCTGAA - - - CCGGAGGACAGGGAG
Hem1ΔD21:
A E K L T I L N - R G Q G V
aa 13
aa 25
C
Nap1/Hem1 KO #7
Nap1/Hem1 KO #7 + EGFP-Hem1ΔD21
Nap1/Hem1 KO #8
Nap1/Hem1 KO #8 + EGFP-Hem1ΔD21
130 kDa
55 kDa
EGFP
Tubulin

#### Figure S5

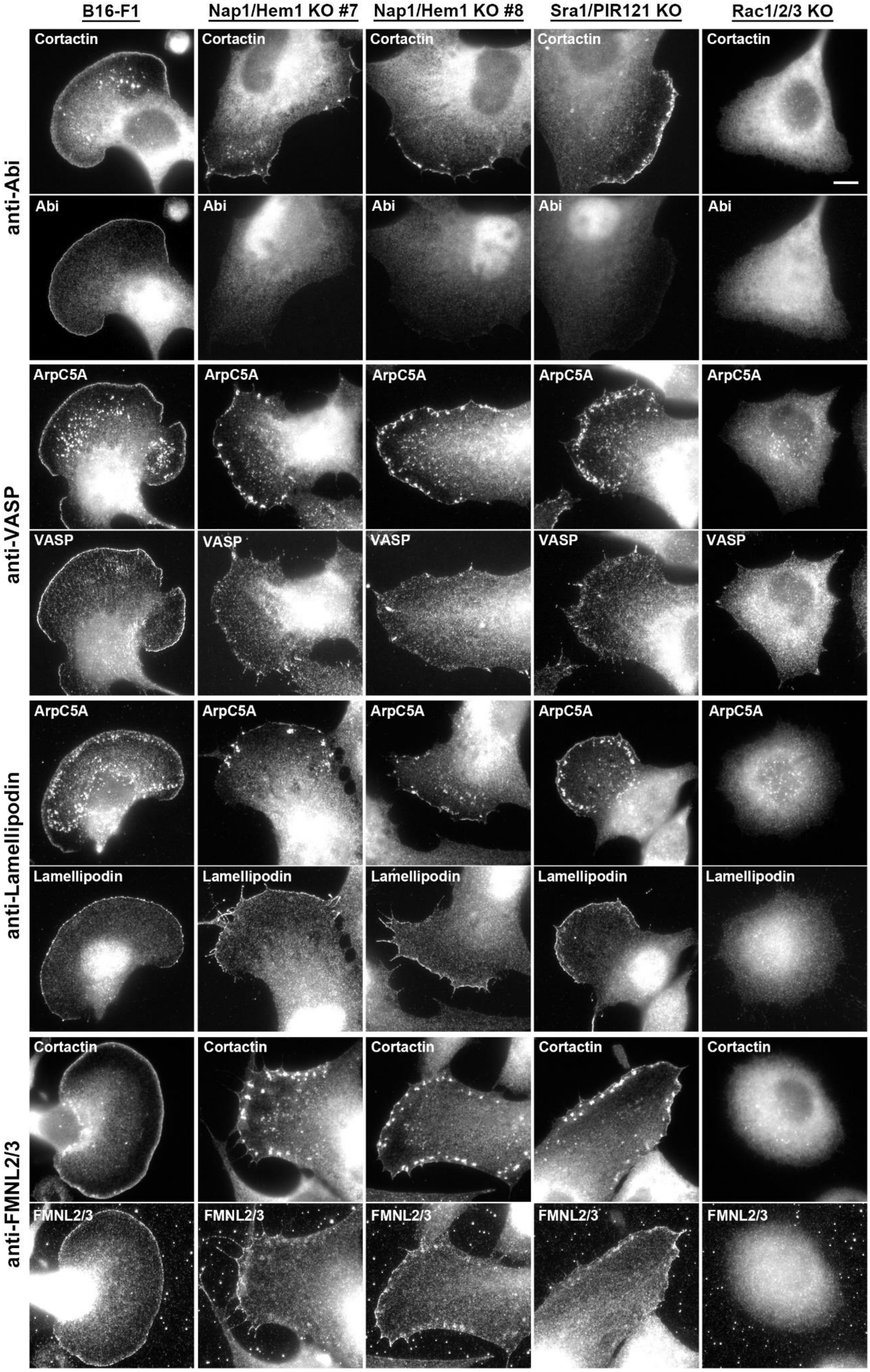
B16-F1
Nap1/Hem1 KO #7
Nap1/Hem1 KO #8
Sra1/PIR121 KO
Rac1/2/3 KO
anti-Abi
Cortactin
Abi
anti-VASP
ArpC5A
VASP
anti-Lamellipodin
ArpC5A
Lamellipodin
anti-FMNL2/3
Cortactin
FMNL2/3

#### Figure S6

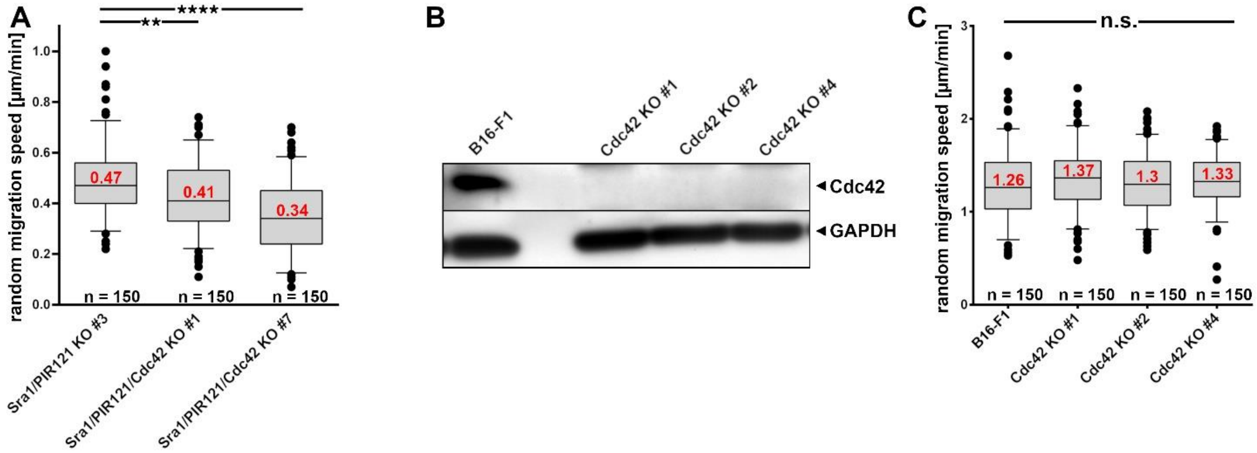
A
B
C
random migration speed [µm/min]
0.47
0.41
0.34
n = 150
n = 150
n = 150
Sra1/PIR121 KO #3
Sra1/PIR121/Cdc42 KO #1
Sra1/PIR121/Cdc42 KO #7
**
****
B16-F1
Cdc42 KO #1
Cdc42 KO #2
Cdc42 KO #4
Cdc42
GAPDH
n.s.
1.26
1.37
1.3
1.33
n = 150
n = 150
n = 150
n = 150

#### Figure S7

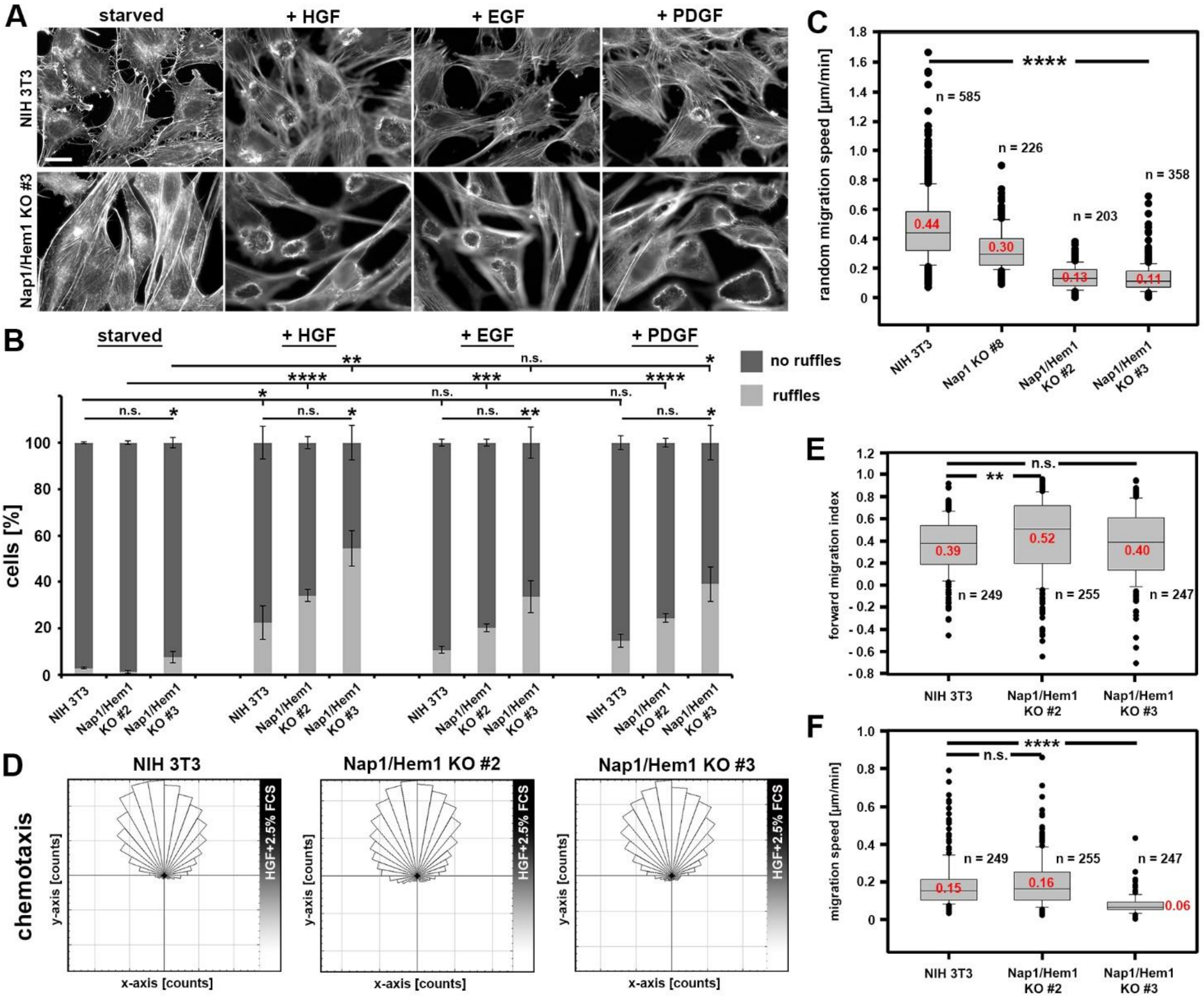

A
starved
+ HGF
+ EGF
+ PDGF
NIH 3T3
Nap1/Hem1 KO #3
B
no ruffles
ruffles
cells [%]
NIH 3T3
Nap1/Hem1 KO #2
Nap1/Hem1 KO #3
C
random migration speed [µm/min]
n = 585
n = 226
n = 203
n = 358
0.44
0.30
0.13
0.11
NIH 3T3
Nap1 KO #8
Nap1/Hem1 KO #2
Nap1/Hem1 KO #3
D
chemotaxis
NIH 3T3
Nap1/Hem1 KO #2
Nap1/Hem1 KO #3
HGF+2.5% FCS
y-axis [counts]
x-axis [counts]
E
forward migration index
0.39
0.52
0.40
n = 249
n = 255
n = 247
F
migration speed [µm/min]
0.15
0.16
0.06
n = 249
n = 255
n = 247
